# Full bandwidth electrophysiology of seizures and epileptiform activity enabled by flexible graphene micro-transistor depth neural probes

**DOI:** 10.1101/2021.09.17.460765

**Authors:** Andrea Bonaccini Calia, Eduard Masvidal-Codina, Trevor M. Smith, Nathan Schäfer, Daman Rathore, Elisa Rodríguez-Lucas, Xavi Illa, Jose M. De la Cruz, Elena Del Corro, Elisabet Prats-Alfonso, Damià Viana, Jessica Bousquet, Clement Hébert, Javier Martínez-Aguilar, Justin R. Sperling, Matthew Drummond, Arnab Halder, Abbie Dodd, Katharine Barr, Sinead Savage, Jordina Fornell, Jordi Sort, Christoph Guger, Rosa Villa, Kostas Kostarelos, Rob Wykes, Anton Guimerà-Brunet, Jose A. Garrido

## Abstract

Mapping the entire frequency bandwidth of neuronal oscillations in the brain is of paramount importance for understanding physiological and pathological states. The ability to record simultaneously infraslow activity (<0.1 Hz) and higher frequencies (0.1-600 Hz) using the same recording electrode would particularly benefit epilepsy research. However, commonly used metal microelectrode technology is not well suited for recording infraslow activity. Here we use flexible graphene depth neural probes (gDNP), consisting of a linear array of graphene microtransistors, to concurrently record infraslow and high frequency neuronal activity in awake rodents. We show that gDNPs can reliably record and map with high spatial resolution seizures, post-ictal spreading depolarisation, and high frequency epileptic activity through cortical laminae to the CA1 layer of the hippocampus in a mouse model of chemically-induced seizures. We demonstrate functionality of chronically implanted devices over 10 weeks by recording with high fidelity spontaneous spike-wave discharges and associated infraslow activity in a rat model of absence epilepsy. Altogether, our work highlights the suitability of this technology for *in vivo* electrophysiology research, in particular, to examine the contributions of infraslow activity to seizure initiation and termination.

## MAIN TEXT

In recent years there has been a resurgence of interest in fluctuations in brain activity occurring at <0.1 Hz, commonly referred to as infraslow activity (ISA)^1^. Several pathological brain states including stroke, traumatic brain injury and migraine with aura are associated with ISA, which can manifest as a slow wave of depolarisation through brain tissue^2^. Interestingly, in the case of epilepsy both fast activity, at hundreds of Hertz (Hz) or higher, and infraslow activity (ISA), at less than 0.5 Hz can be associated with seizures and epileptiform activity^3^. Moreover, seizure generation has been hypothesised to be generated by a coupled dynamical system in which there are fast and slow processes^4^. However, the relationship between these two types of brain activity is poorly understood. A limitation in studying ISA, either independently or concurrently with higher frequency activity is the lack of appropriate tools to record it electrographically *in vivo* with high spatiotemporal fidelity. Experimentally ISA is usually recorded using solution-filled glass micropipettes with Ag/AgCl wires which limits the spatial resolution to just a few-point measurements. A further issue with glass pipettes is that they are not practical for long-term chronic recordings in awake animals and are incompatible with clinical use. To enhance the spatial resolution and long-term recordings, microelectrode grids can be used, however this is not optimal since they suffer from polarization-induced drift and signal attenuation causing distortion of the measured signal^5^. Consequently, research investigating the relationship between ISA and higher frequency activity, either in normal or pathological brain, is hampered by a lack of appropriate technology.

An alternative to commonly used passive electrodes are field-effect transistors (FETs), which are active transducers offering significant advantages in electrophysiology, in particular the capability to monitor infraslow signals^6^. Among the few FET technologies that have been validated for *in vivo* electrophysiology, graphene-based technology is particularly attractive because of the combination of special properties of this material, including chemical and electrochemical inertness, high electrical mobility, biocompatibility, as well as a facile integration into flexible and ultrathin substrates^7^. Recent reports demonstrate the potential of graphene solution-gated field-effect transistors (gSGFETs) for neural interfacing^8,9^. A first proof-of-concept demonstration using epicortical gSGFET arrays for mapping chemically-induced ISA in anesthetized rats has been reported^6^. To advance this technology further, we have developed implantable flexible graphene depth neural probes (gDNP) capable of recording localised full bandwidth neuronal activity, through cortical columns and sub-cortical structures in preclinical rodent models of induced seizures and chronic epilepsy.

Here, we demonstrate a wafer-scale microtechnology process to fabricate gDNPs consisting of a linear array of graphene micro-transistors imbedded in a polymeric flexible substrate. In order to penetrate through the mouse cortex and reach the hippocampus without buckling, we adapted an insertion protocol that uses silk-fibroin (SF)^10,11^ to temporary stiffen flexible gDNPs. We validate experimentally the ability to detect electrophysiological biomarkers of epileptiform activity, including high frequency oscillations (HFOs)^12,13^ comparable to conventional microelectrodes and we highlight the suitability of graphene transistor technology to record concurrently additional biomarkers in the infraslow frequency range^14^. These include DC shifts preceding seizure onset^15,16^ and post-seizure spreading depressions^17^, accentuating the potential of this technology to gain mechanistic insight into the involvement of infraslow activity associated with seizures *in vivo* in awake brain.

## Results

### Microfabrication and characterization of the gDNPs

A graphene-based SGFET is a three terminal device in which single layer CVD graphene is used as the channel material in contact with the drain and the source terminals. Graphene is the sensing part of the device directly exposed to the neural tissue. The current in the graphene channel can be modulated or pinned by a third terminal given by a reference electrode (gate) which is also in contact with the conducting neural tissue (Fig. 1a). Thus, variations in the electrical potential in the tissue can be transduced into variations of the channel current; this transduction mechanism has been shown to offer a very versatile sensing platform for electrophysiology^18,19^. The flexible gDNP is a linear array of 14 recording transistors, each with an active area of 60×60 μm, and a pitch of 100 μm. The probe’s tip design consists of a polyimide shank of 200 μm width and 1.6 mm length (Fig. 1b). gDNPs are fabricated on a 10 μm thick flexible polyimide (PI) substrate using a wafer-scale microfabrication process previously reported^9^ (see Methods). A two-level metallization strategy, with metal levels interconnected using via-holes (Fig. 1b), reduces track resistance and improves sensor performance. To characterize gDNPs in saline solution we measure simultaneously the drain-source current (I_DS_) versus the applied gate-source voltage (V_GS_) for all the transistors on the shank with a fixed drain-source voltage (V_DS_) using customized electronics (see Methods). The transistor sensitivity is a function of its transconductance (g_m_), defined as g_m_ = dI_DS_/dV_GS_, which is directly linked to the ability of the gSGFET to amplify recorded signals. gSGFETs exhibit very high g_m_ values due to the large capacitive coupling at the graphene-electrolyte interface and to the very high charge carrier mobility of graphene^20^. Fig. 1c shows the transfer curves as well as the normalized transconductance (g_m_/V_DS_), for all 14 gSGFETs of an exemplary gDNP device. The small dispersion of the charge neutrality point (CNP), defined as the value of V_GS_ where the I_DS_ reach its minimum, attests for the homogeneity of the gDNP. This is crucial for its operation *in vivo* because all transistors share a common source terminal. Furthermore, g_m_ shows a very stable response in a wide applied gate frequency range (up to 5 kHz), both in the hole regime, V_GS_ < V_CNP_, and in the electron regime, V_GS_ >V_CNP_ (Fig. 1d). Such constant frequency response is important for a proper calibration of the recorded signals^21^. The detection limit of the sensors is evaluated by means of the effective gate noise (V_RMS_) integrated between 1 Hz and 2 kHz, with averaged values between 25-30 μV for all fabricated gDNPs (see Supplementary Fig. S2).

**Fig. 1.**
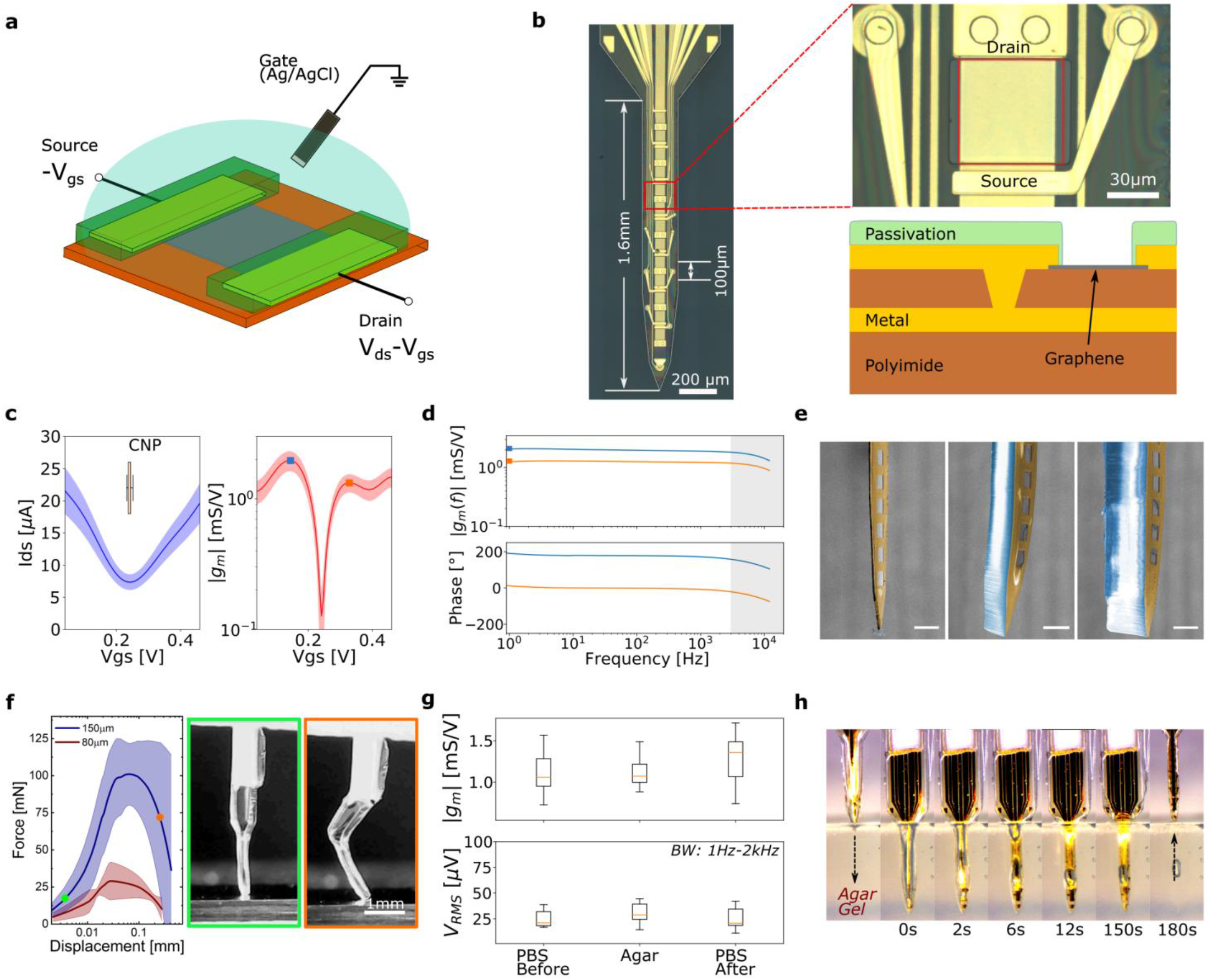
Flexible graphene Depth Neural Probe (gDNP) technology and characterization. **a**, Schematic of a graphene solution-gated field-effect transistor (gSGFET) and biasing. Vgs: gate-source voltage, V_ds_: drain-source voltage. **b** Optical microscope image of a gDNP containing 14 transistors with a pitch of 100 μm on a 200 μm wide polyimide shank. Right: blown up image of one gSGFET; the red contour highlights the graphene sensing area (60×60μm) of the transistor. The schematic of the cross section of one transistor shows the interconnected metal tracks strategy to reduce the shank width of the gDNP. **c-d**, Electrical characterisations of all 14 gSGFET on a gDNP in a 150 mM saline solution (V_ds_ = 50 mV). **c**, Mean values with standard deviation (shaded colours) of drain-source current (I_ds_) and transconductance (g_m_) versus Vgs. **d**, Transconductance spectroscopy of the gSGFET bias at the point of maximum g_m_ in the electron (Vgs > V_CNP_, orange line) and hole regime (V_gs_ < V_CNP_, blue line). Squares dots are the values of the g_m_ as measured in steady-state mode. The decay observed in the grey shaded areas is due to the filtering of the interfacing electronics. **e,** Coloured SEM images of the gDNP; uncoated (left), back coated with ~80 μm (middle) and with ~150 μm thick silk-fibroin (right) (scale bar=100 μm). **f**, Mechanical assessment: averaged compression force vs displacement for the gDNP coated with two SF thicknesses (coloured areas are standard deviations, n=10 trials); the optical images correspond to two different conditions of the experiment. **g**, Functionality assessment: maximum normalized transconductance (g_m_/V) values and averaged V_RMS_ electronic noise level, of all gSGFET on a device measured in a PBS solution, inserted and measured in agarose gel brain model, and measured in the PBS solution after removal from agarose gel. **h**, Image sequence of a SF-coated gDNP at different time points during degradation in 0.6% agarose gel brain model.

### Stiffening of the flexible gDNP using silk-fibroin

gDNPs are highly flexible, compared to traditional rigid depth electrodes, and although flexibility is highly advantageous once inserted into the tissue, this provides a challenge during insertion. To insert these probes we temporarily stiffen the gDNP using silk-fibroin (SF)^10,11^. Compared to other natural biopolymers, SF offers excellent mechanical properties, extremely good biocompatibility, biodegradability, and the versatility of structural readjustments^22,23,24^. Further, the byproducts of the SF degradation by enzymes (e.g. proteases) have low antigenicity and non-inflammatory characteristics^25,26^. The stiffening technique (see Methods and Fig. S3) consists of a moulding process in which the gDNP is back-coated with cured SF, allowing the preparation of a rigid, straight shank with a defined shape and thickness. We tuned the thickness of the SF by controlling the mould’s trench depth, achieving two typical thicknesses of 80±10 μm and 150±12 μm, as shown in the scanning electron microscopy image of Fig. 1e. Mechanical assessment of the SF coated gDNP was performed using a buckling test, in which the perpendicularly positioned probes were driven against a flat and hard surface (Fig. 1f). An initial linear increase in force is observed for both coating thicknesses tested, while the probes remained straight before buckling (green box in Fig. 1f). Continued application of force results in buckling and bending (orange box in Fig. 1f), characterized by a peak in the force-displacement curve. The obtained peak forces, 101±21 mN for the 150 μm thick SF and 29±13 mN for 80 μm thick SF, are in good agreement with the previously reported values of peak forces of similar SF-coated neural probes^11,27^. In order to evaluate the effect of the stiffening and insertion procedures on the device performance, we electrically characterized the gDNPs before and after the SF stiffening process, as well as before and after insertion and removal from an agarose gel brain model. Fig. 1g shows the averaged values of the normalized g_m_ as well as the effective gate noise (V_RMS_) of all 14 transistors on a gDNP, confirming that neither the stiffening process nor the insertion in an agarose brain model impair gDNP performance in terms of transconductance nor noise. Video frames of a SF-coated gDNP inserted in an agarose brain model (Fig. 1h) show the fast water absorption (<10 s) of SF after complete insertion (insertion speed: 400 μm/s) as well as the collapsing of the SF in small residue beads which often stays in the solution for a longer period of time. As observed from the gDNP after removal from agar gel (180 s), SF is completely delaminated from the polymeric shank, therefore making the SF coated probes suitable for single-time insertion.

### Awake *in vivo* full bandwidth recording with gDNPs

We assessed full bandwidth recording capability by implanting a gDNP into awake, head-fixed mice. The electrophysiological signal measured by the graphene transistors was acquired with a g.RAPHENE system (g.tec medical engineering GmbH) that enables simultaneous recording in two frequency bands with different gains preventing amplifier saturation (Fig. 2a, Methods). gDNPs were implanted in the right hemisphere visual cortex (V1) and lowered until the tip reached hippocampal tissue. Baseline activity was recorded for (10-20 min). To induce network discharges and synchronicity of neuronal bursting 200nl of 4-AP (50mM), a selective blocker of Kv1 potassium channels^28,29^ was focally injected into cortex adjacent to the gDNP (Fig. 2b).

**Fig. 2.**
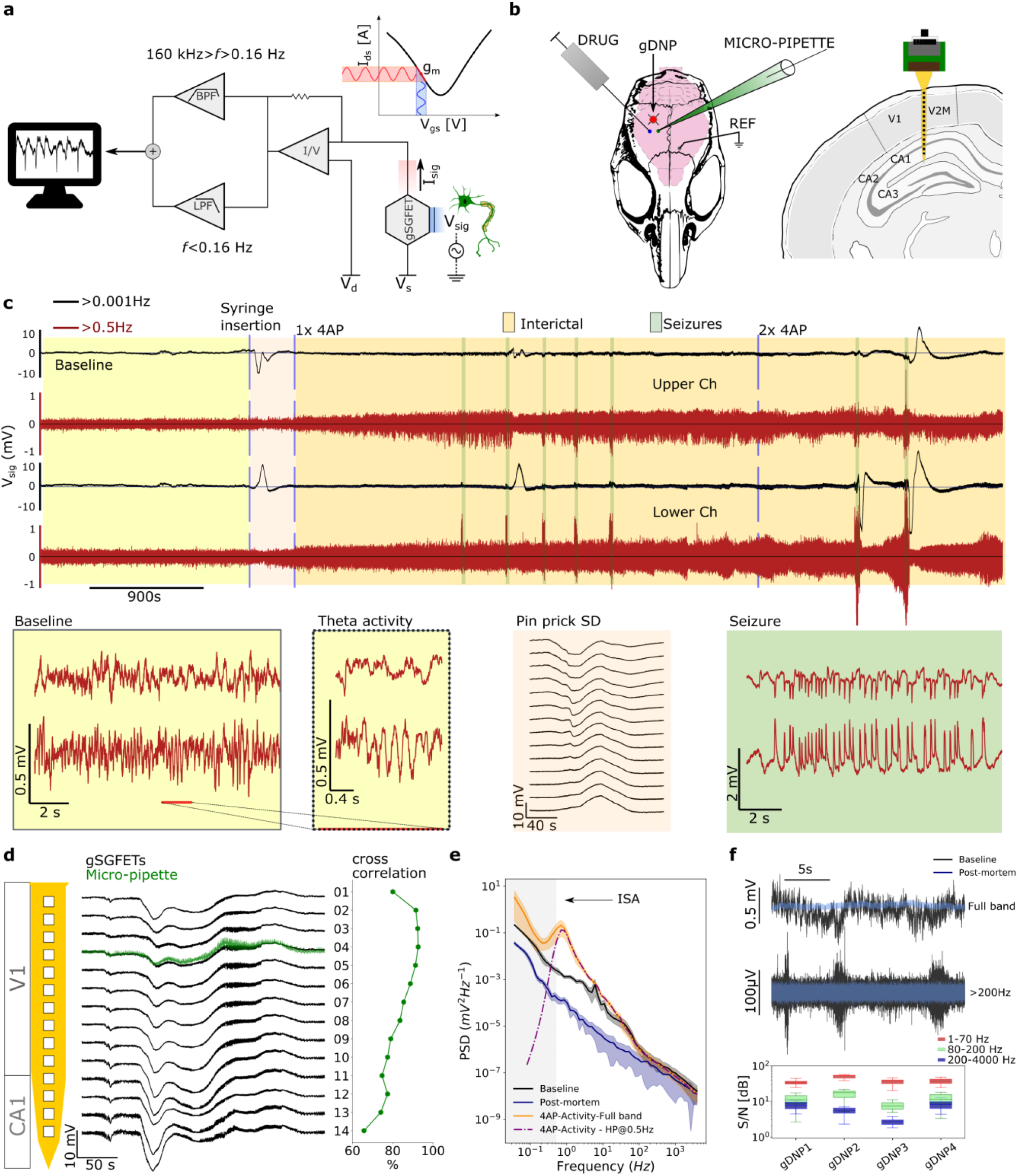
Validation of *in vivo* full bandwidth recording capabilities of gDNPs. **a,** Schematic of the recording setup and concept of a graphene transistor as a transducer for neural recording. **b,** Schematic of a mouse skull with location on the brain of the gDNP, the glass micropipette and the needle to inject the chemoconvulsant drugs. Right, coronal view of the mouse brain with the localisation of the gDNP. **c**, Long electrophysiological recording (120 mins) of two channels on the gDNP array (top: visual cortex, bottom: hippocampus). The full bandwidth (f > 0.001 Hz, black) signal and the HP filtered signal >0.5 Hz (dark red). Baseline activity, pin-prick SD, increased neuronal activity after 4-AP injection and seizures, some of them followed by a post-ictal SD. Below, different events in higher resolution: baseline (yellow) revealing theta activity in the lowest channel; profile visualization with recording from all 14 transistors following pinprick SD (beige); seizure activity shown for the uppermost and lowest channels of the gDNP (green). **d**, The recording shows the full bandwidth epileptiform activity followed by an SD event from all 14 transistors (black). The superimposed green recording corresponds to the signal measured with the glass micropipette. The subfigure shows the low frequency cross-correlation (< 5 Hz), between the micro-pipette and all transistors on the shank. **e,** Averaged PSD over the electrophysiological recordings of all transistors during baseline, epileptiform activity, same activity HP filtered at 0.5 Hz (purple dashed-line) and post-mortem. The grey area highlights the low frequency part (<0.5 Hz) usually cut-off with conventional AC-coupled recordings. **f,** Comparison of a baseline activity (black) and a post-mortem (blue) in one channel of the gDNP (top: full band, bottom: HP>200 Hz). Lowest plot shows SNR evaluation for 4 *in vivo* experiments performed with 4 different gDNPs. The SNR is calculated for different bands (LFP:1-70 Hz, high frequency: 80-200 Hz and very high frequency: 200-4000 Hz) and is the averaged SNR for all channels on each gDNP.

#### Full bandwidth recordings

Fig. 2c displays 2 hours of an electrophysiological recording session (only the uppermost and the lowest channels of the implanted gDNP displayed); the complete data set is shown in Supplementary Fig. S6. The ability of the graphene transistors to have long and stable full-bandwidth recordings without the need for electronic off-set readjustments contrasts to the limitation of DC-coupled passive electrodes^30^. The black lines correspond to the full bandwidth signal (HP > 0.001 Hz) and the red lines to the signal high-pass filtered above 0.5 Hz (which is the expected signal recorded by AC-coupled electrodes)^31^. The coloured regions correspond to different experimental conditions during the recording: baseline (yellow), needle-induced pin-prick SD^32^ (pink), interictal activity^14^ (orange) induced by chemoconvulsant drugs, and seizures (green & see Supplementary Fig. S7). During baseline recording in Fig. 2c, lower channels exhibit theta activity, correlated with animal movement, indicating that the gDNP reached the hippocampus, confirmed post-hoc by histological analysis of fixed brain sections (see Supplementary Fig. S8). After injection of 4-AP epileptiform spiking evolved and five seizures (over 60 minutes) were elicited in this example, one of which was followed by a post-ictal SD. A second cortical injection of 4-AP induced two additional seizures both followed by post-ictal SDs that were detected first in the hippocampus. In 5 different mice injected with 4-AP, an average number of 7 ± 3 seizures were recorded in 60 min post drug injection. In this chemoconvulsant model SDs could be observed initially either in superficial cortical layers or, the hippocampus (Fig. 2c).

#### Validation of infraslow activity recordings with glass micropipette

The fidelity of recorded ISA activity was validated by simultaneous recordings using a solution-filled glass micropipette, which is considered the gold-standard for ISA recordings. Fig. 2d shows the full bandwidth recording obtained with the gDNP (black lines) and the micropipette (green line) after injection of 4-AP. Both recordings reveal DC shifts preceding the seizure and a high amplitude ISA occurring after the seizure. The ISA deflection measured by the gDNP has a similar shape, magnitude and temporal duration as the signal recorded by the glass micropipette. A cross-correlation analysis (signal filtered <5 Hz) of the signal recorded by the glass micropipette and the 14 gDNP transistors demonstrates a very high correlation (above 90%) for ch03 and ch04 located at the same cortical depth as the micropipette.

#### Assessment of the detection limits of gDNP

Post-mortem recordings were acquired to characterize the electrical noise level of the gDNP in the activity-free brain state and, consequently, to quantify the detection limit of the gDNP. Fig. 2e shows the averaged power spectral density (PSD) calculated using the recordings of all channels in a gDNP, obtained from different brain states (baseline, after injection of 4-AP, and post-mortem). Compared to the baseline PSD, the large amplitude of the PSD at low frequencies (< 1 Hz) after 4-AP injection is an indication of the interictal and infraslow activity in the brain. The dash-dot line in Fig. 2e corresponds to the activity recorded after injection of 4-AP, but with a typical HP filter of 0.5Hz, found in many AC-coupled recording systems, thus revealing the loss of ISA signal (grey area). In order to assess the detection limit in the conventional frequency bands (>1Hz), the recordings of the baseline were directly compared with post-mortem recordings. For instance, applying a digital filter (> 200 Hz) and comparing post-mortem with baseline validates the ability of gDNP to record spontaneous high-frequency activity (> 200 Hz) arising from groups of neurons in a non-pathological brain state. Beyond this qualitative comparison, we have calculated the signal-to-noise ratio (SNR) in three different bands, 1-70Hz (red), 80-200Hz (green), and 200-4000 Hz (blue) for different gDNPs implanted in four animals. SNR is calculated as root-mean square (RMS) amplitude ratio of the baseline and post-mortem recordings, filtered in the three different bands (see Methods). These results show that the gDNPs are able to record typical electrophysiological signal bandwidths with SNR ratios higher than 1 dB.

#### Interictal activity and HFOs

Fig. 3a shows interictal activity and associated HFOs (>80 Hz) ^12,13^ recorded by three of the transistors of a gDNP, each located at a different depth in the mouse brain. Filtering between 80 – 600 Hz (red curves in Fig. 3a) reveals layer-specific bursting of HFOs and sharp wave ripples, during inter-ictal spikes with characteristic oscillations of 200-300 Hz and 400-600 Hz in the cortical and hippocampal channels respectively^33^ (Supplementary Fig. S9); entrained inter-ictal epileptiform activity was found in all channels before each seizure. Two different pro-convulsive drugs (4-AP or picrotoxin PTX) were used to induce and evaluate epileptic activity. Fig. 3b illustrates characteristic examples of sharp wave ripples and HFOs induced by 4-AP and by PTX recorded by the lowest channel of the gDNP (hippocampal CA1 region). The HFO and ripple traces shown in Fig. 3b exhibit high-frequency tones up to 600 Hz. The filtered traces (>200 Hz) are compared to the original traces (full-bandwidth) for verification of ripples. The advantage of the gDNPs to monitor concurrent infraslow and high-frequency activity, is illustrated in Fig. 3c, which shows a post-ictal spreading depression (SD) arising from the hippocampus. The layer-dependent silencing of the neural activity by the hippocampal SD is represented in Fig. 3d in terms of activity variation [%]. The right plot in Fig. 3d shows the layer-dependent amplitude of the SD and the following hyperpolarization along the vertical profile (Supplementary Fig. S10 and Methods), revealing that the silencing of the neural activity in the hippocampus is correlated with the amplitude and subsequent hyperpolarization wave of the SD. Silencing of neural activity in the hippocampus by the SD is visualized with more clarity in Fig. 3e, where the spectrograms for the upper and lowest channel are compared. Fig. 3f shows more details of the seizure event recorded at three different depths, from the visual cortex to the hippocampus, comparing the signal in full band (black curves), HP filtered >0.5Hz (red curves), and LP filtered < 0.5Hz (green curves).

**Fig. 3.**
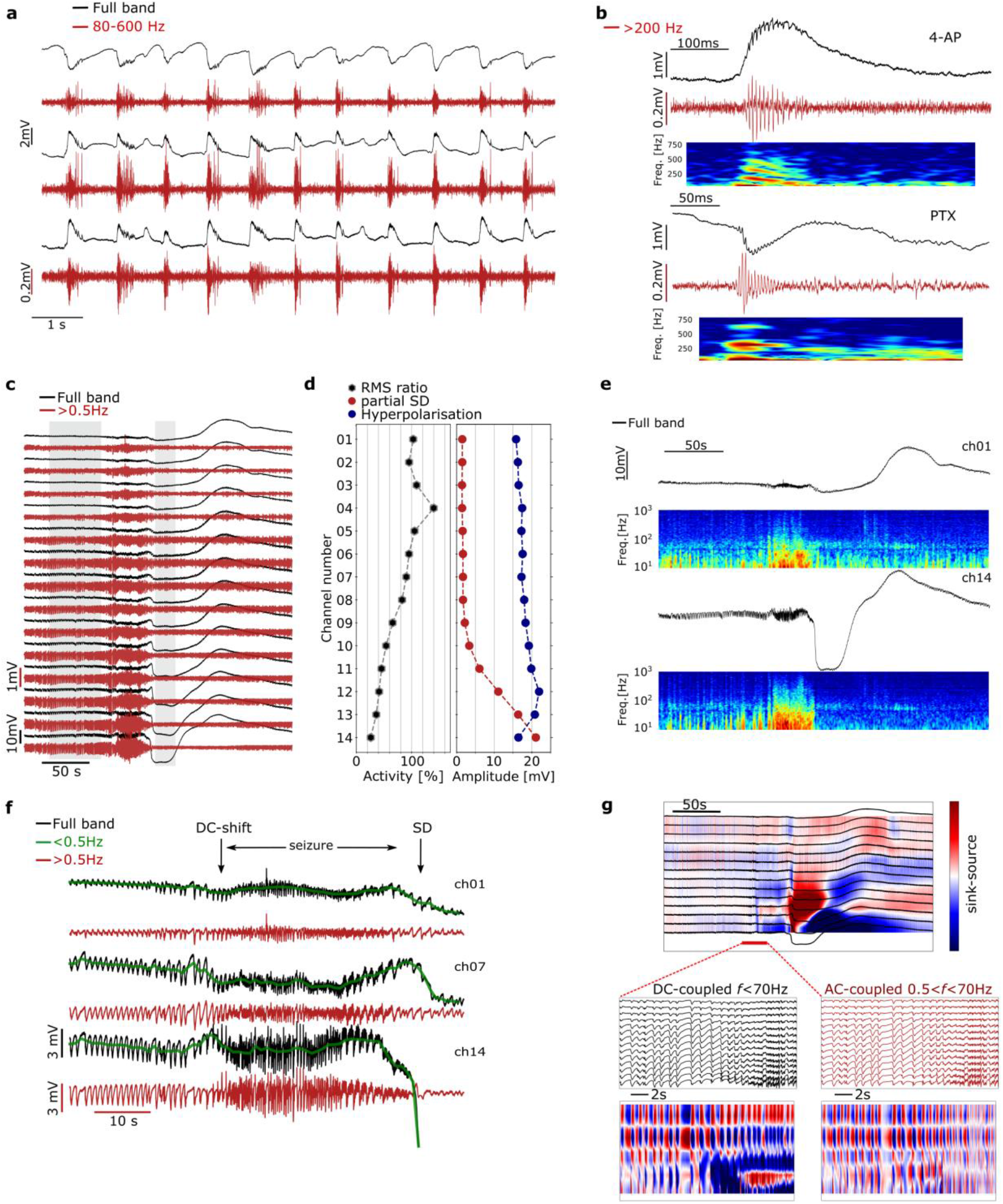
Electrophysiological recording of characteristic epilepsy biomarkers. **a,** Interictal activity in three different channels of a gDNP (ch01, ch07, ch14) (red curves BPF 80 Hz – 600 Hz). **b**, Sharp-wave ripples and HFO recorded in the hippocampus induced by 4-AP and PTX (full band: black, HP>200 Hz: red); the figure also shows the corresponding spectrograms (range 10-800 Hz). **c,** Electrophysiological full band recordings (black curves) and HP filtered at 0.5 Hz (red curves) from the cortex (top channel) to hippocampus (bottom channel) illustrating a post-ictal spreading depression (SD) arising from the hippocampus after a seizure event. **d**, Neural activity variation (before seizure and during SD, grey area in (**c**)) and amplitudes of the SD and hyperpolarisation waves concurrent with the seizure across the vertical profile. Neural activity variation is calculated for each channel by comparing activity before and during the SD (grey shaded areas in panel (**c**)). **e**, Hippocampal neural silencing during the SD is illustrated by the spectrograms (range 10-1000 Hz) of the uppermost and lowest channels of the gDNP (ch01, ch14). **f**, Three channels (ch01, ch07, ch14) showing full band recording (black) and band-pass filtered (> 0.5 Hz, red) of a seizure followed by a hippocampal SD; in green the low-freq. component of the recording (< 0.5 Hz) overlapped to the full bandwidth signal, showing an ictal DC shift associated with a seizure, and followed by the SD. **g,** Current-source density (CSD) analysis of the low frequency activity (< 10 Hz), showing source and sinks in the hippocampus during the SD. Below: blow up of the pre-ictal to seizure transition (< 70 Hz), showing dipoles in the different layers of the cortex and hippocampus. The two graphs correspond to the CSD analysis performed with (left) and without (right) the contribution of low frequency components.

#### Current-source density analysis using infraslow activity

Ictal DC shifts can be associated with seizures^16,34^ but are usually removed from recordings due to the requirement for high-pass filtering applied to conventional AC-coupled electrodes^31^. We are able to record these using gDNPs and observe DC shifts immediately prior to seizure onset, with their amplitude (0.5-3mV) and polarity related to cortical layer (Fig. 3f). The inversion of DC shifts can be used to identify current sources and sinks through the cortical laminae. Current-source density (CSD) analysis is a technique to identify source activation in a variety of focal neurological disorders including epilepsy^35^. CSD analysis (see Methods) of the data in Fig. 3c reveals a large net ionic sink in the hippocampal extracellular space after the seizure, followed by a large source at the beginning of the hyperpolarization wave (Fig. 3g). Enlarging the seizure onset region, 4 sink and source regions are identified through the laminae profile. CSD analysis computed without the infraslow components (0.5Hz<f<70Hz) fails to report the ionic sinks preceding and during the seizure in the bottom layers (Fig. 3g, AC-coupled panel), illustrating the importance of using full-bandwidth recordings for CSD analysis to avoid misinterpretation of the extracellular potential sinks and sources (Supplementary Fig. S11 illustrates additional examples of CSD analyses).

### Chronic functional validation and biocompatibility assessment

Finally, we discuss the stability and chronic functionality of gDNP, defined by the ability to maintain a suitable signal-to-noise ratio recording spontaneous epileptiform activity over time. We implanted gDNPs in the right-hemisphere somatosensory cortex of WAG-Rij rats (n=4), a rodent model of absence epilepsy^36^, and obtained chronic full-bandwidth recordings over a 10-week period Fig. 4a. WAG-Rij rats exhibit frequent spontaneous spike-and-wave discharges (SWDs), a characteristic thalamocortical oscillation between 8-10 Hz^36^. Implanted animals were connected 1-2 times per week for tethered recordings (using a commutator to enable free movement of the rats see Fig. 4a).

**Fig. 4.**
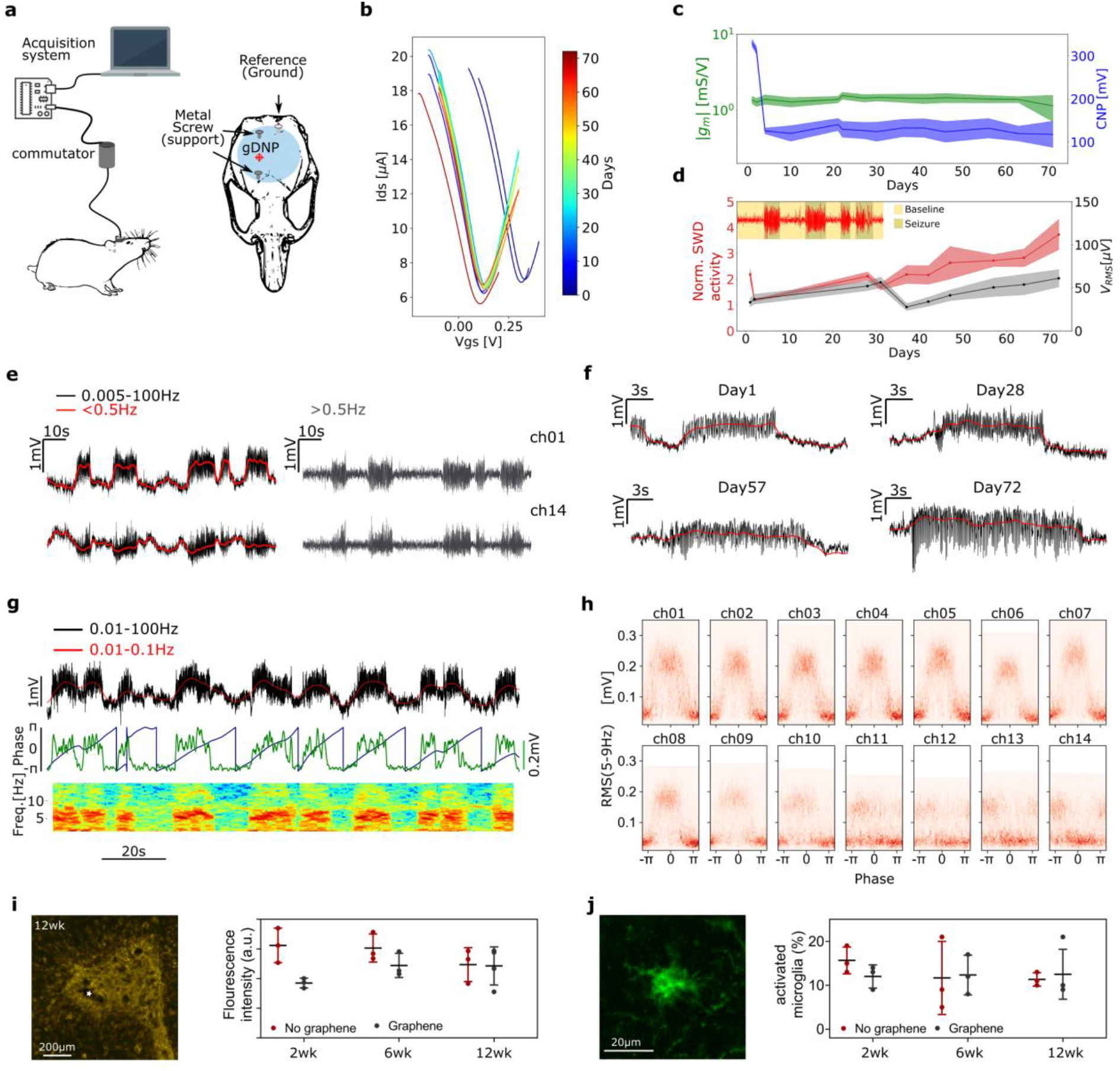
Chronic assessment of stability, functionality and biocompatibility of gDNPs. **a,** Left: Cartoon of a WAG-Rij rat tethered via a commutator and the electronics for the data acquisition. Right: Schematic of the rat skull with the approximate location of the gDNP. **b-d,** Stability and quality of the recordings. **b,** Averaged transfer curves of all transistors of the gDNP over the implantation time. **c,** Averaged values of the maximum transconductance g_m_ (green) and the charge neutrality point (CNP, blue) of the transistors transfer curves. **d,** Normalized SWD activity over implantation time, calculated as the ratio of the SWD activity and baseline activity. Inset: example of SWDs events (red), highlighting the periods considered baseline and SWD. The graph also shows the average V_RMS_ noise (black) of all channels over the implantation period. **e,** SWD in the uppermost and lowest channels of the gDNP; ISA component (red<0.5 Hz) overlapped to the 0.005-100 Hz signal (black); in grey the same signal filtered above 0.5 Hz. **f,** Illustrative SWD events measured by the same channel at days 1, 28, 57, and 72. **g,** Typical recording of one of the transistors located in the rat cortex showing SWDs events concurrent with ISA. Phase of the ISA (blue) and power of the SWD activity (green). The spectrogram (range 1-14 Hz) shows the increase in power during the SWDs. **h**, Density distribution for all channels evaluated in a long recording (1600s). y-axes correspond to the RMS (5-9 Hz) associated with the SWD and the x-axes represent the phase of the ISA (red represents higher density); shows a clear correlation between ISA phase and SWD power with a phase inversion in the deeper cortex layers. **i-j**, Chronic biocompatibility assessment of SF-coated gDNPs implanted in Sprague-Dawley rats (n=20) by monitoring inflammatory markers. **i**, Fluorescent image of GFAP, as a measure of positive astrocyte cells, in the area of insertion at 12 weeks post implantation (the star mark shows the insertion point). Time evolution (2, 6, 12 weeks) of the fluorescence intensity (calculated at 150μm from the probe sites) obtained for flexible gDNP with graphene (grey), and gDNP without graphene (red). **j**, Example of microglial activation (fluorescence image). Time evolution (2, 6, 12 weeks) of activated microglia in the vicinity of the implanted probes (in an area of 0.7 mm^2^) obtained for the two types of probes (graphene, no-graphene).

#### Stability of implanted transistors

Transistor curves were measured in each recording session to assess device stability, changes at the device/tissue interface and importantly, to permit selection of an optimal Vgs to maximise SNR; a feature possible with active sensor devices^19^. Fig. 4b shows the averaged transfer curves of a gDNP measured over 10 weeks (see also Supplementary Fig. S12). The stability of the transistors’ performance is illustrated in Fig. 4c, which depicts the position of the CNP and the maximum value of g_m_ over the implantation period. The averaged g_m_ value remains approximately constant over the whole study. Since g_m_ is directly related to the tissue/graphene interfacial capacitance and to the carrier mobility in graphene, the g_m_ stability strongly suggests little or no variation of these two parameters. CNP (Fig. 4c) shows a significant shift (200 mV) during the few first days after implantation, which then remains stable for the rest of the experiment. We tentatively attribute the initial shift to the adsorption of negatively charged species, which reduce the intrinsic p-type doping of the graphene transistors^20^.

#### Long-term functionality of gDNPs

Long-term functionality of the gDNP was assessed by evaluating the quality of the recorded signals over the implantation period using two parameters, normalized SWD power amplitude and the transistor noise (V_RMS_). Normalized SWD power amplitude is defined as the RMS (calculated in the frequency band 1-500 Hz) of the SWD activity normalized by the baseline activity (non-SWD periods, see inset of Fig. 4d and Supplementary Fig. S13). V_RMS_ over time is extracted by averaging the RMS values at very high frequency (500-2000 Hz) in the non-SWDs periods of the recording, where we expect the power of neural activity to be closer to the transistor’s noise (see Fig. 2f). For further details, refer to Methods and Fig. S12. In Fig. 4d the variation of these two parameters over time is shown, demonstrating the ability of the implanted devices to monitor seizure activity with high fidelity during the whole implantation period. The slight increase in the normalized SWD activity could result from a strengthened coupling between neural tissue and the gDNP or from an increase in the seizure power and duration as the animal ages^37^.

#### Correlation between SWDs and infraslow activity

The WAG Rij rat chronic model of absence epilepsy offers the possibility to investigate correlations between infraslow activity and SWD events. Because of the full-bandwidth capability of the gDNP, we were able to observe DC shifts (< 0.5 Hz) associated with SWDs, and a correlation between the amplitude and polarity of the DC shift with the depth of the neocortex layers. Fig. 4e shows the uppermost and lowest channel with opposite associated DC shifts during each SWDs. In this experiment we observed a positive DC shift (~1.5 mV) in the superficial layers L1, L2/L3, and L5 to a lesser degree, and a negative DC shift in the lowest cortical layers (Supplementary Fig. S14). The latter is not observed after application of a HP-filter (>0.5 Hz) as typically used with conventional AC-coupled microelectrodes (Fig. 4e, grey). The observed correlation between the SWDs and ISA, and layer-dependent polarity of the DC shift persist over the implantation period. Representative seizure events showing SWDs and DC shifts over time are shown in Fig. 4f.

We calculated the phase of the ISA (limited to the 0.01-0.1 Hz band) and the power of the neural activity associated to the SWDs (5-9 Hz). Fig. 4g displays one of the upper channels together with its spectrogram (range 1-14 Hz). The distribution of the ISA phase with the power amplitude of the SWDs is plotted for each channel (Fig. 4h), showing a clear inversion in the lower layer of the neocortex (see Methods and Supplementary Fig. S15). This correlation between ISA and SWD was also observed in the other implanted WAG-Rij rats (n=4, Supplementary Fig. S16). gDNPs are therefore a promising electrophysiology tool to gain further understanding of the influence of thalamocortical oscillations in SWD generation^38^.

#### Chronic biocompatibility of implanted gDNP

In addition to the chronic functional validation of the gDNP, we conducted an extensive chronic biocompatibility study to assess any potential neuro-inflammation caused by the invasive nature of the penetrating neural probes, the presence of CVD graphene, or by the release of SF following implantation. gDNPs with or without graphene at the recording sites were implanted in adult, male Sprague-Dawley rats (n=20). Histological and immunohistochemical studies were conducted at 2, 6, and 12 weeks’ post-implantation and compared to the contralateral hemisphere, without device implantation (see Methods and Supplementary Fig. S17, Fig. S18). Fig. 4i shows a fluorescence image of GFAP immunostaining (positive marker for astrocyte cells) in the area of insertion 12 weeks post implantation (brain sections at 800 μm from pia). There is no significant increase in the presence of astrocyte cells (typically associated with inflammation) in the area directly surrounding the probe site, when the “graphene” gDNP is compared with the “no-graphene” gDNP (Fig. 4i, right graph). Moreover, no significant difference was observed at 2, 6, and 12 weeks’ post-implantation. Further, the microglial activation state, assessed by morphological analysis of cells stained for ionized calcium binding adaptor molecule 1 (Iba-1), showed no significant increase in the abundance of activated microglia present in the area surrounding the implantation site (Fig. 4j). Additional immunohistochemical analysis showed no sign of an inflammatory response over the 12-week period for either device used (Supplementary Fig. S17). Altogether, the chronic biocompatibility study indicates that gDNPs are suitable for chronic implantation, inducing no significant damage nor neuroinflammatory response.

### Outlook

In this work, we demonstrate the capability of flexible depth neural probes based on linear arrays of graphene microtransistors to map the entire frequency bandwidth, simultaneously recording infraslow activity and high frequency neural oscillations through the cortex laminae to upper hippocampal layers. Measuring the full bandwidth of neuronal activity in the brain with high spatiotemporal resolution will advance our understanding of the brain, in health and disease. While SD and seizures can occur concurrently, the inter-layer dynamics and the effect of SD on epileptic activity across the cortical laminae and deeper regions of the brain remain largely unknown. gDNPs allowed us to reveal associations between infraslow activity(including spreading depolarizations^17,39^ and small pre-seizure DC shifts^15,16,40^) and higher frequency activity (including SWDs^38^ and HFOs^12,13^) in rodent models of drug-induced seizures and chronic epilepsy^36^. Together with validation of chronic functionality of implanted gDNPs and their biocompatibility, this work underlines very distinct advantages of this technology for *in vivo* epilepsy preclinical research. We envision that uptake of this technology will aid mechanistic understanding of seizure initiation and termination and gain insight into the nature of post-seizure spreading depolarisations, recently implicated in sudden unexplained death in epilepsy (SUDEP)^41,42^. Clinical development could result in depth probes capable of simultaneous high-quality wide-band (DC to HFO) recordings, from multiple brain regions, during pre-surgical monitoring improving identification of seizure onset zones (SOZ) and ultimately surgical outcome^14^. Beyond epilepsy research, this technology is expected to advance our understanding of neurological diseases and could easily be applied to study disorders associated with infraslow activity including traumatic brain injury, stroke and migraine^2^.

## Methods

### Graphene growth, transfer and characterization

Graphene was grown by chemical vapor deposition (CVD) on copper foil (Alfa Aesar Coated). Prior to growth, copper foil was electropolished for 5 min in a H2O (1 L) + H3PO4 (0.5 L) + ethanol (0.5 L) + isopropanol (0.1 L) and urea (10 g) solution^43^. The CVD reactor consists of a tubular three zone oven with a quartz tube (1600×60 mm). After loading Cu foil, an annealing step (1h) was performed, prior to growth, at 1015°C under a 400 sccm Argon flow at 100 mbar. This was followed by the growth step (15 min growth step), at 12 mbar under a gas mix of 1000 sccm Argon, 200 sccm hydrogen, and 2 sccm methane. Transfer of the graphene from copper foil to polyimide was achieved using a wet-etching chemical method. A supporting poly(methyl methacrylate) PMMA 950K A4 was spin-coated on the graphene/copper foil and left to dry for 12 h. Back side graphene is removed with a 10% HNO3 solution. Subsequently, the sample was laid on the etchant solution composed of FeCl3/ HCl (0.5M/2M) to remove copper for at least 6 h. Next the graphene/PMMA stack was rinsed in DI water multiple times before transfer onto the polyimide-coated wafer. The wafer was dried for 30 min at 40 °C on a hot plate, and annealed in a vacuum hotplate by increasing the temperature gradually up to 180 °C. Finally, the PMMA was removed in acetone and isopropanol. CVD-grown graphene was characterized by Raman spectroscopy using a Witec spectrograph equipped with a 488 nm laser excitation line. To assess the quality of the graphene film (see Supplementary Fig. S1) Raman maps of 30 x 30 μm^2^ (with 1 μm resolution) were acquired using a 50x objective and the 600 g nm^-1^ grating; laser power was kept below 1.5 mW to avoid sample heating.

### Microfabrication of flexible gDNPs

Flexible neural probes were fabricated using standard microelectronic fabrication technology on a rigid 4-inch sacrificial Si/SiO2 wafer. A 10 μm thick polyimide (PI-2611, HD MicroSystems) layer was spin-coated and cured at 350 °C in a N2 atmosphere. To reduce the shank width of the depth neural probes, we used a two-metal level strategy in which the metal tracks, separated by PI, are interconnected by via-holes. After evaporation and definition (via lift-off) of a first metal layer of Ti/Au (10/100 nm), a 3 μm-thick PI layer was spin-coated and cured. A protective mask of Al (200nm) was used to etch the second PI layer by oxygen plasma and form the via-holes. On top of the via-holes a second metal layer of Ti/Au (10/100 nm) was applied to interconnect the two metal layers. CVD graphene was transferred onto the patterned wafer as described in the section above. After removing the PMMA protection layer, the graphene active areas were defined by means of an oxygen-based reactive ion etching. A sandwich-like contact strategy was used to improve the contact at the drain and source terminals; the used top metal structure was Ni/Au (20/200nm). For passivation, a 2μm-thick chemically resistant polymer is deposited (SU8-2005 MicroChem) with open windows in the channel region. Finally, the gDNP structure was defined in a deep reactive ion etching process using the thick AZ9260 positive photoresist (Clariant) as an etching mask. The polyimide probes were then released from the SiO2 wafer and placed in a zero insertion force connector in order to interface our custom electronics.

### Characterization of gDNPs in saline

The graphene SGFETs on the neural probes were characterized in PBS solution (150mM). Drain to source currents (Ids) were measured varying the gate–source voltage (Vgs), versus a Ag/AgCl reference electrode which was set to ground. Steady state was ensured by acquiring the current only after its time derivative was below a threshold (10^-7^ A s ^-1^). The detection limit of the graphene SGFET were assessed by measuring the power spectral density of the DC current at each polarization point Vgs. Integrating the PSD over the frequencies of interest (1Hz-2kHz) and using the transconductance allowed us to calculate the root-mean-square gate voltage noise VRMS. The noise measurement was performed in a Faraday cage, with DC-batteries powering the amplifiers, in order to reduce any 50 Hz coupling or pick-up noise. Additionally, the frequency response of the device’s transconductance was measured by applying a sum of sinusoidal signals at the electrolyte solution through the reference electrode and by measuring the modulation of the drain current. The acquired signals were split into two bands, low frequencies (≈ 0–10 Hz), at which drain–source current was simultaneously acquired for all transistors in an array, and higher frequencies (10 Hz–30 kHz), at which each transistor was recorded individually.

### Back coating of gDNP with silk fibroin

Temporary stiffening of the gDNPs with silk-fibroin is achieved using a micro-structured PDMS mould with the shape of the neural probes. To fabricate the moulds, PDMS is cast on a standard 4-inch silicon wafer with pre-patterned 100 μm and 200 μm thick SU8 (SU8-2050) epoxy resin. The back-coating procedure is as follows: first, the probe is placed in the mould trench filled previously with water, with the transistor side facing down. Through surface tension the probe self-aligns in the mould. After evaporation of the water, silk fibroin (Sigma Aldrich, Silk, Fibroin Solution 50 mg/mL) was applied via a syringe to the mould’s trench. We double coat the shank in drying intervals of 20 minutes and then slowly increase the temperature on a hotplate to 80 °C, leaving the SF curing for 1h 30 minutes. By increasing the duration of the water annealing step we managed to have a delayed dissolution time compared to SF cured at room temperature. After curing, the coated probe can be easily removed from the PDMS mould (see Supplementary Fig. S3). In all presented *in vivo* experiments we implanted the flexible gDNPs with a 150 μm thick SF back-coating.

### Assessment of mechanical properties of the stiffened gDNPs

Standard compression tests against a hard silicon (Si) substrate were performed to assess the mechanical properties of our SF-coated probes. Buckling experiments were carried out in a UMIS nanoindenter from Fischer-Cripps Laboratories. A custom clamp was fabricated to fix the probes at the end of the indenter shaft that, in turn, was connected to the actuator and load cell. Buckling tests were carried out at a loading rate of 8.8 mN/s. Once the indenter detected noticeable buckling the test was automatically stopped. The maximum applied load that the indenter can apply is 500 mN. Applied force vs displacement was measured until the probe started buckling and eventually broke down. We additionally measured the Young’s modulus of the SF cured at 80 °C by means of nano-indentation tests. SF was drop casted on a 2×2cm2 Si chip and cured. The indentation measurements were performed using a NHT2 Nanoindentation Tester from Anton-Paar equipped with a Berkovich pyramidal-shaped diamond tip. A maximum applied load of 5 mN was applied with a loading segment of 30 s followed by a load holding segment of 10 s and an unloading segment of 30 s. The hardness and reduced Young’s modulus are reported as an average value of at least twenty indentations, performed on top of each sample (in the central region). Young’s modulus values in the range of 10 GPa where measured for 80 °C cured SF (see Supplementary Fig. S4).

### Electronics for in vivo recordings with gDNPs

The experimental setup used to perform the *in vivo* recordings provides Vs and Vd bias control and current-to-voltage conversion for up to 16 channels (g.RAPHENE, g.tec medical engineering GmbH, Austria). The instrumentation splits the recorded signals into two bands with different gains: low-pass filtered (LPF, < 0.16 Hz, 10^4^ gain) and band-pass filtered (BPF, 0.16 Hz < f < 160 kHz, 10^6^ gain). Two custom Simulink models were used: i) to perform the transfer curve of the microtransistors once inserted and at the end of the experiment; ii) to set the Vs and Vd bias and acquire the recorded signals. Signals were sampled at 9.6 kHz and at 19kHz depending on the type of experiment (see Supplementary Table1).

### Ethical approval and animal handling for acute and chronic experiments

Animal experiments were conducted in accordance with the United Kingdom Animal (Scientific Procedures) Act 1986, and approved by the Home Office (license PPL70-13691). C57BL/ mice were bred (2-4 month old males), while WAG rats were imported (Charles river, used 6-9 months of age). Animals were housed on 12 h/12 h dark/light cycle, and food and water were given *ad libitum*. Prior to headbar sugery, animals were group housed, but after this, animals were individually housed.

### Acute preparation surgeries for headbar attachment and craniotomy

For both surgeries, aseptic techniques were used with mice anaesthetized using isoflurane (988-3245, Henry Schein, U.S.A.) and placed in a stereotaxic frame (David Kopf Instruments Ltd., U.S.A.). Viscotears applied (Bausch + Lomb, U.S.A.) and pain relief, which consisted of sub-cutaneous Bupenorphine (0.5 mg / Kg;Ceva, France) and Metacam (15 mg /Kg; Boehringer Ingelheim, Germany), were injected. Saline was administered just before recovery or every 45 mins depending on the length of surgery. To apply the headbars for the Neurotar system the skin on the top of the head was cut to expose the skull. The skull was cleaned and dried, which enabled drilling (RA1 008, Henry Schein, U.S.A.) of a small hole in the left hand visual cortex for a metal support screw (00-96X3-32, Plastics One, U.S.A.). Using vetbond (1469SB, 3M, U.S.A.), the headplate (Model 9, Neurotar, Finland) was firmly attached and strengthened using dental cement before Kwik-cast (KWIK-CAST, W.P.I., U.K.) covered the exposed skull. Mice were checked daily to ensure recovery. After at least 5-days of recovery, habituation was performed by placing the mouse in the Neurotar frame for increasing periods of time (15-60 mins) over several days. On the day of recording, a craniotomy was performed. Under Isoflurane anaesthesia, with administration of pain medication and intra-muscular Dexamethasone (1 mg / Kg; intra-muscular; 7247104; MSD Animal Health, U.S.A.), two areas were exposed. A large (2×2mm) craniotomy over somatosensory and visual cortex on the right-hand side and a small drill hole over the motor cortex on the left hand side. Cold Cortex buffered saline was continually applied to the craniotomies. After completion, exposed dura was covered with Cortex buffered saline, sterilised slygard (~200 μm thickness), and a kwik-cast layer. After ~2-hours post-recovery, the animal was moved to the Neurotar frame and the craniotomies were exposed by removal of the kwickcast and sylguard. The gDNP was carefully connected to a PCB and lowered using a micromanipulator to just above the dura over the visual cortex. The dura was gently pierced using either micro-dissection scissors or a 26-gauge needle and gDNP lowered ~2 mm into the brain. A reference wire (Ag/AgCl_2_) was placed in the ipsilateral motor cortex and g.tec hardware (see *Electronics for in vivo recordings with gDNPs*) used to perform a DC characterisation curve to determine the optimal Vgs and initiate recordings. Chemoconvulsant was injected into the brain using a Nanofil injection system (W.P.I., U.K.). At the end of the experiment, sodium pentobarbital was administered intra-peritoneally.

### Recording with solution-filled glass micropipette

Borosilicate capillary tubes (OD: 1.50mm, ID: 0.86mm, Warner Instruments) were pulled using a horizontal puller (Sutter instruments P-97, resistance of 3-5 MOhm) and filled with artificial Cerebral Spinal Fluid (NaCl: 119mM, KCl: 2.5mM, CaCl2: 2.5mM, MgSO4: 1.3mM, NaH2PO4: 1.25mM, NaHCO3: 25mM, Glucose: 10mM) and attached to an Axon instruments CV-7B head stage. A micro-manipulator (MM3301, WPI) was used to position the pipette above the cortical surface before insertion approximately 400μm into the cortex. The head-stage was provided with the same reference as the gDNP, a chlorinated silver wire touching the ipsilateral motor cortex. The headstage was connected to a Multiclamp 700B amplifier (Axon Instruments) operating in current clamp mode. Analogue-Digital Conversion and TTL pulse delivery for temporal synchronisation was achieved using the Micro1401 MkII (CED, Cambridge, U.K.). Data was acquired using WINEDR sampling at 20 kHz with a 4 kHz Bessel filter.

### Chronic preparation surgery and recording

First, the gDNP was fibroin coated, as described above, to aid insertion. The rat was anaesthetised to a surgical depth using Isoflurane. After placement in a stereotaxic frame, Viscotears were applied and pain medication, which consisted of Bupenorphine (0.15 mg / Kg; sub-cutaneous; Ceva, France) and Metacam (4.5 mg /Kg; sub-cutaneous; Boehringer Ingelheim, Germany), was applied. The skull was cleaned and dried. Small burr holes (~1 mm) were drilled at four positions: 1) Somatosensory cortex for gDNP (since the perioral somatosensory cortex is the focal area for SWDs ^44^); 2) contralateral cerebellum for a reference Ag/AgCl wire held in place by a nylon screw; 3) Motor cortex, ipsilateral, for a support screw; and 4) Visual cortex, ipsilateral, for a support screw. The metal screws were inserted and provided structural support for the dental cement. Next, the gDNP and the reference wire were inserted, and a DC characterisation curve confirmed that the transistors were performing optimally. Dental cement, mixed with vetbond, was applied around the PCB for support. Animals were weighed daily and their physiology was monitored to ensure a full recovery. For recording, animals were anaesthesia-free and moving, with the PCB-interface on the head connected to an Omnetics cable (A79635, Omnetics, U.S.A.) that interfaced with the g.tec recording hardware as described above. After ~5-minutes for settling, a DC characterisation curve was recorded to allow accurate calibration of the gSGFETs. A script calculated the optimal Vgs based on the transfer curves. Recordings were performed for ~10-60 minutes twice a week for 10 weeks. After recording, the Omnetics wire was disconnected and a protective cap was applied.

### Data analysis

All electrophysiological data were analysed using Python 3.7 packages (Matplotlib, Numpy, Neo and Elephant) and the custom library PhyREC (https://github.com/aguimera/PhyREC). The conversion of the recorded current signals (LPF and BPF) to a voltage signal was performed by summation of the two signals and interpolation in the in vivo/chronic measured transfer curve of the corresponding gSGFET. The transfer curves were always measured at the beginning and end of every recording to ensure that no significant variations were present and to detect any malfunctioning transistor. Moreover, all recordings presented in the manuscript have been calibrated with the nearest-recorded transfer curve to ensure high fidelity in the voltage-converted signals. The SNR shown in Fig. 2f was evaluated by the ratio of the RMS mean value over 25s of recording of baseline (spontaneous activity) and post-mortem (no activity). The signal is BP filtered in three different bands corresponding to the LFP activity (1-70 Hz), high frequency (80-200 Hz) and very high frequency activity (200-4000 Hz). RMS values were calculated with a sliding window of 500ms for the 1-70 Hz band and with a sliding window of 10ms for the other two bands. For each BP filtered signal the mean RMS ratio is calculated and averaged for all 14 channels in the gDNP. SNR for the different bands is evaluated from a total of 4 in-vivo experiments with 4 different gDNPs (Fig. 2f). SNR is expressed in dB (20*ln(*RMS*(*S*)/*RMS*(*N*))). The silencing of neuronal activity shown in Fig. 3d, was extracted using the AC-coupled recording (HP >0.5 Hz). Then the RMS values of the pre-ictal phase (calculated with a sliding window of 1s) are averaged over 50s time. Similar analysis was performed for the time during the SD (15s, shaded areas in Fig. 3d). The ratio of the two averaged RMS values corresponds to the neuronal activity variation [%] (before and during the SD). The amplitude of the hippocampal SD and hyperpolarization wave in Fig. 3d, is evaluated using the recording LP filtered in the infra-slow regime (<0.5 Hz) and re-sampled at 3 Hz (instead of 9.6 kHz used in the original recording). The zero of the voltage was set using the mean value of the signal 50s before the pre-ictal phase (see Supplementary Fig. S10), the minimum and maximum values for each channel were extracted (corresponding to the SD and the hyperpolarisation amplitude respectively). Current source density analysis applied to the low-frequency part of the potential (LFP), was calculated with the python open source Elephant library (Elephant electrophysiology analysis toolkit) using the class “Current Source Density analysis (CSD)” and the method 1D – StandardCSD was chosen for the linear gDNP array. A homogeneous conductivity of the neural tissue of σ=0.3 S/m across the different layers was used for the calculations. The ISA concurrency with the SWD shown in the histogram in Fig. 4h was evaluated by performing the Hilbert transform to extract the phase of the ISA (0.01-0.1 Hz) and the RMS between 5-9 Hz (typical bandwidth for the SWD). Supplementary Fig. S15 shows in more detail the dependency of ISA phase and SWD amplitude.

### Chronic biocompatibility study

#### Device manufacture and sterilization

Two types of flexible gDNP where fabricated for the immunohistochemical study: One with graphene and one without graphene following the fabrication steps described above section (*Fabrication of gDNPs*). In the devices gDNP without graphene the graphene instead of being defined by RIE, was etched away. By doing so, we make sure that all the fabrication steps are equal for both, gDNP with and gDNP without graphene. For comparison to rigid devices currently available on the market, iridium Neuronexus electrodes (A1×32-Poly2-5mm-50s-177) with a thickness of 15μm and length 5mm were implanted. Devices were sterilised individually with ethylene oxide, using an Anprolene AN-74i sterilizer, performed according to manufacturer’s instructions.

#### Surgical implantation of devices

Adult male Sprague-Dawley rats (230-280g) were used for this study (Charles River, England). All animals were kept in individually ventilated cages (Techniplast, GR1800) in groups of 3-4, housed at a constant ambient temperature of 21 ± 2°C and humidity of 40–50%, on a 12-h light, 12-h dark cycle. All rats were given free access to diet and water. Experimental procedures were conducted in compliance with the Animal welfare act 1998, with approval of the Home Office and local animal welfare ethical review body (AWERB). Animals were anaesthetized with Isoflurane (2-3%) throughout surgery, and depth of anaesthesia was monitored with the toe pinch reflex test. Animals were fixed to a stereotaxic frame (Kopf, model 900LS), and body temperature was maintained with a thermal blanket. A small craniotomy (~3mm) was made with a micro drill (WPI, OmniDrill35) above the somatosensory cortex, the dura was excised and one of three depth probe devices were implanted; i) graphene device, ii) no graphene device, or iii) Neuronexus device, at coordinates relative to bregma; anteroposterior (AP): 0mm, dorsoventral (DV): +3.5mm, and mediolateral: −1.5mm. The craniotomy site was sealed with Kwik Sil (WPI), secured with dental cement, the skin was sutured closed, and anaesthetic was withdrawn, with saline (20ml/kg) and buprenorphine (0.03mg/kg in saline) given subcutaneously to replace lost fluids and reduce post-operative pain.

#### Tissue collection and processing

Animals were culled at 2, 6 or 12 weeks’ post-implantation dependent on the analysis to performed. Tissue was taken either for immunohistochemical analysis of cells related to inflammatory processes, or for cytokine analysis of inflammatory markers. For a further description of the techniques used, and analysis see Supplementary Information.

#### Histology

Animals were perfused with 4% PFA to fix tissue. Axial plane brain sections were cut at 50μm thickness with a vibrotome (Leica, VT1200). Sections at an approximate electrode site depth of 0.8mm were selected for staining. Sections were stained free-floating for two markers; i) ionized calcium binding adaptor molecule 1 (Iba1) to quantify microglial population, or ii) Glial fibrillary acidic protein (GFAP) staining to assess astrocyte presence (see Supplementary Methods for details of immunohistochemistry). Slides were imaged with a Leica SP8 confocal microscope with a 10x objective lens. Laser power and digital gain was kept consistent across imaging sessions. A single optical section of the tissue surrounding the probe sight was taken within the middle portion of the section as to avoid edge effects. Details of the analysis of histology is provided in Supplementary Methods section.

#### Enzyme-Linked ImmunoSorbent Assay (ELISA) Protocol

For ELISA, animals were culled by rising concentration of CO2. Brain tissue was extracted, snap frozen in liquid nitrogen, and stored at −80 °C until further use. Brain tissue was lysed by addition of NP-40 lysis buffer (150 mM NaCl, 50 mM Tric-Cl, 1% Nonidet P40 substitute, Fluka, pH adjusted to 7.4) containing protease and phosphatase inhibitor (Halt™ Protease and Phosphatase Inhibitor Cocktail, ThermoFisher Scientific) followed by mechanical disruption of the tissue (TissueLyser LT, Qiagen). Samples were centrifuged at 5000RPM for 10 minutes, and the supernatant stored at 4 °C until further use. A bead-based multiplex ELISA kit was run, which included markers interleukin-1a (IL-1a), interleukin-1beta (IL-1b), interleukin-17 alpha (IL-17a), and interleukin-33 (IL-33) (Cat. No. 740401, Biolegend). The standard instructions for the kit were used, with protein loaded at a fixed volume of 15 μL. After incubation, beads were run on the BD FACSVerse flow cytometer, and the data analysed using LEGENDplexTM Data Analysis software.

#### Statistical analysis

For histological staining, all data sets are n=3, with the exception of 2-week graphene, which is n=4, and 12 weeks Neuronexus probes which are n=2, due to an inability to locate the probe location in histological sections for one animal implanted. For ELISA testing, gDNP with and without graphene hemisphere data sets are n=3 or 4 at all timepoints, while contralateral hemispheres were combined, giving n=7.

## Funding

This work has been funded by the European Union’s Horizon 2020 research and innovation programme under Grant Agreement No 785219 (Graphene Flagship). The ICN2 is supported by the Severo Ochoa Centres of Excellence programme, funded by the Spanish Research Agency (AEI, grant no. SEV-2017-0706), and by the CERCA Programme / Generalitat de Catalunya. A.B.C. is supported by the International PhD Programme La Caixa - Severo Ochoa (Programa Internacional de Becas “la Caixa”-Severo Ochoa). This work has made use of the Spanish ICTS Network MICRONANOFABS partially supported by MICINN and the ICTS ‘NANBIOSIS’, more specifically by the Micro-NanoTechnology Unit of the CIBER in Bioengineering, Biomaterials and Nanomedicine (CIBER-BBN) at the IMB-CNM. This work is within the project FIS2017-85787-R funded by the “Ministerio de Ciencia, Innovatión y Universidades *“* of Spain, the “Agencia Estatal de Investigación (AEI)” and the “Fondo Europeo de Desarrollo Regional (FEDER/UE)”. A.B.C. acknowledges that this work has been done in the framework of the PhD in Electrical and Telecommunication Engineering at the Universitat Autònoma de Barcelona.

R.W. is funded by a Senior Research Fellowship awarded by the Worshipful Company of Pewterers

D. R. is a Biotechnology and Biological Sciences Research Council (BBSRC) LIDo sponsored PhD student.

The authors would like to thank Prof. Matthew Walker and Prof. Louis Lemieux (UCL Queen Square Institute of Neurology) for their comments on the manuscript.

## Author contributions

A.B.C. carried out most of the fabrication and characterization of the gDNPs, contributed to the design and performance of the in vivo experiments, analysed the data and wrote the manuscript. E.M.C. contributed to the design and planning of the in vivo experiments and support to the SD analysis of the in vivo data. R.W., T.M.S. performed the in vivo experiments. D.R., contributed to the in vivo experiments and DC-coupled recordings with the glass micropipette. N.S., E.R.L., X.I. and J.M.D.C. contributed to the fabrication and characterization of the gDNPs. E.D.C., J.B. and C.H., contributed to the growth, transfer and characterisation of CVD graphene used in the gDNPs. E.P.A., A.H. and E.R.L., contributed to the optimisation of the silk-fibroin stiffening protocol of the gDNPs. J.M.A., contributed to the fabrication of the custom electronic instrumentation and development of a Python-based user interface. D.V. contributed to the python scripts and technical discussions. J.R.S. reviewed the manuscript. J.F. and J.S. contributed to the mechanical assessment of the silk-fibroin and the silk-fibroin back-coated gDNPs. M.D. performed all surgeries for the biocompatibility study. A.D. and K.B. contributed to the capture of histological images and image-processing and analysis. S.S and K.B contributed to the preparation and review of the manuscript. A.G.B. contributed in the design and fabrication of the custom electronic instrumentation, development of a custom gSGFET Python library and analysis of the data. R.V., K.K., R.W., A.G.B., and J.A.G. participated in the design of the in vivo experiments and thoroughly reviewed the manuscript. All authors read and reviewed the manuscript.

## Data and materials availability

Raw data from gDNP characterization, electrophysiological recordings as well as biocompatibility assessment will be shared upon request to the corresponding authors.

