## Supplementary Information for "Full bandwidth electrophysiology of seizures and epileptiform activity enabled by flexible graphene micro-transistor depth neural probes"

#### Supplementary Methods

##### Chronic biocompatibility study

**Histology.** At the end of the experimental period (2, 6 or 12 weeks) rats were anaesthetized with Isoflurane and culled via cardiac perfusion with heparinised (10U/ml, Sigma-Aldrich) phosphate buffered saline (PBS), followed by 4% Paraformaldehyde (PFA, Sigma-Aldrich) in PBS. Brains were then post-fixed in 4% PFA in PBS for 24 hours, transferred to 0.01% Sodium Azide PBS (Sigma S-8032) thereafter, and stored at 4°C. Axial plane brain sections were cut at 50μm thickness with a vibrotome (Leica, VT1200). Sections at an approximate electrode site depth of 0.8mm were selected for staining. Sections were stained free-floating for two markers; i) ionized calcium binding adaptor molecule 1 (Iba1) to quantify microglial population, or ii) Glial fibrillary acidic protein (GFAP) staining to assess astrocyte presence. Tissue sections were stained with rabbit anti-Iba1 (1:1000, Wako), and chicken anti-GFAP (1:2000, Abcam ab4674) overnight at 4°C. Sections were incubated with the secondary antibodies; anti-rabbit Alexa Fluor (AF) 560, and anti-chicken AF647 (all 1:500, Thermofisher) for 2 hours at room temperature. Sections were mounted onto slides and coverslips were mounted with Prolong Gold antifade mounting media (Thermofisher). Slides were imaged with a Leica SP8 confocal microscope with a 10x objective lens. Laser power and digital gain was kept consistent across imaging sessions. A single optical section of the tissue surrounding the probe sight was taken within the middle portion of the section as to avoid edge effects. Microglial cells were individually classified into one of four morphologies; Grade 0 (resting/ramified), Grade 1 (de-ramifying/re-ramifying), Grade 2 (activated/amoeboid) or Grade 3 (clustered & activated). Activation was determined as a percentage of total microglial cells which were either Grade 3 or 4.

**Immunohistochemical data analysis.** Raw TIF files were loaded into a custom Python3.7 script (github.com/kebarr/biocompatibility_study). We averaged the data obtained from each section in an animal prior to plotting, so each data point corresponds to a single animal. The values for each section were obtained as described below.

**i) GFAP evaluation.** The fluorescence intensity of the normalised GFAP images was assessed as a function of distance from the probe site. First, the probe site was localised in each image, based on the bright fluorescence around its edges (Supplementary Fig. S17). The region identified as the probe site was used as a base mask for quantifying the fluorescence intensity, and the intensity in the masked area was summed so that it could be later subtracted when calculating the intensity in bands around the probe site. To quantify the intensity within a distance of one pixel from the edge of the probe site, the binary dilation operation was applied to the probe site mask. Then, the intensity within the dilated mask was summed, and the intensity inside the region of the base mask was subtracted, leaving the total intensity within the ring around the probe site. This process was then repeated, always subtracting the total intensity from the previous masked area as we iterated outwards up to 200 pixels from the probe site. To visualise this data, all the curves obtained for each condition were smoothed using a Savitzky-Golay filter, then averaged by the lineplot function in the Python seaborn (version 0.9.0) package, giving one final curve.

**ii) Iba1 evaluation.** IBA1 fluorescence images were used to quantify the microglial activation. First, the cells were segmented based on pixel values exceeding a simple, manually set, threshold. The segmented cells were then labelled using the 'measure.label' function from scikit-image (v 0.16.2), and their region properties were quantified using the 'measure.regionprops' function. One of these properties is called the 'extent,' which compares the area of the bounding box of the labelled object to the area of the object itself. Spindly objects, such as microglia that are not activated, will have a large bounding box for a relatively small object area, leading to a small extent. Rounder objects without long protrusions, such as activated microglia, will occupy a large amount of their bounding box, leading to a larger extent. Hence, this metric can be used to classify whether microglia are activated or not. For an example of how this classification strategy works, please see Supplementary Fig. S18. Manual cross-checking of a number of cell counts was performed to validate the automated classification.

### Results

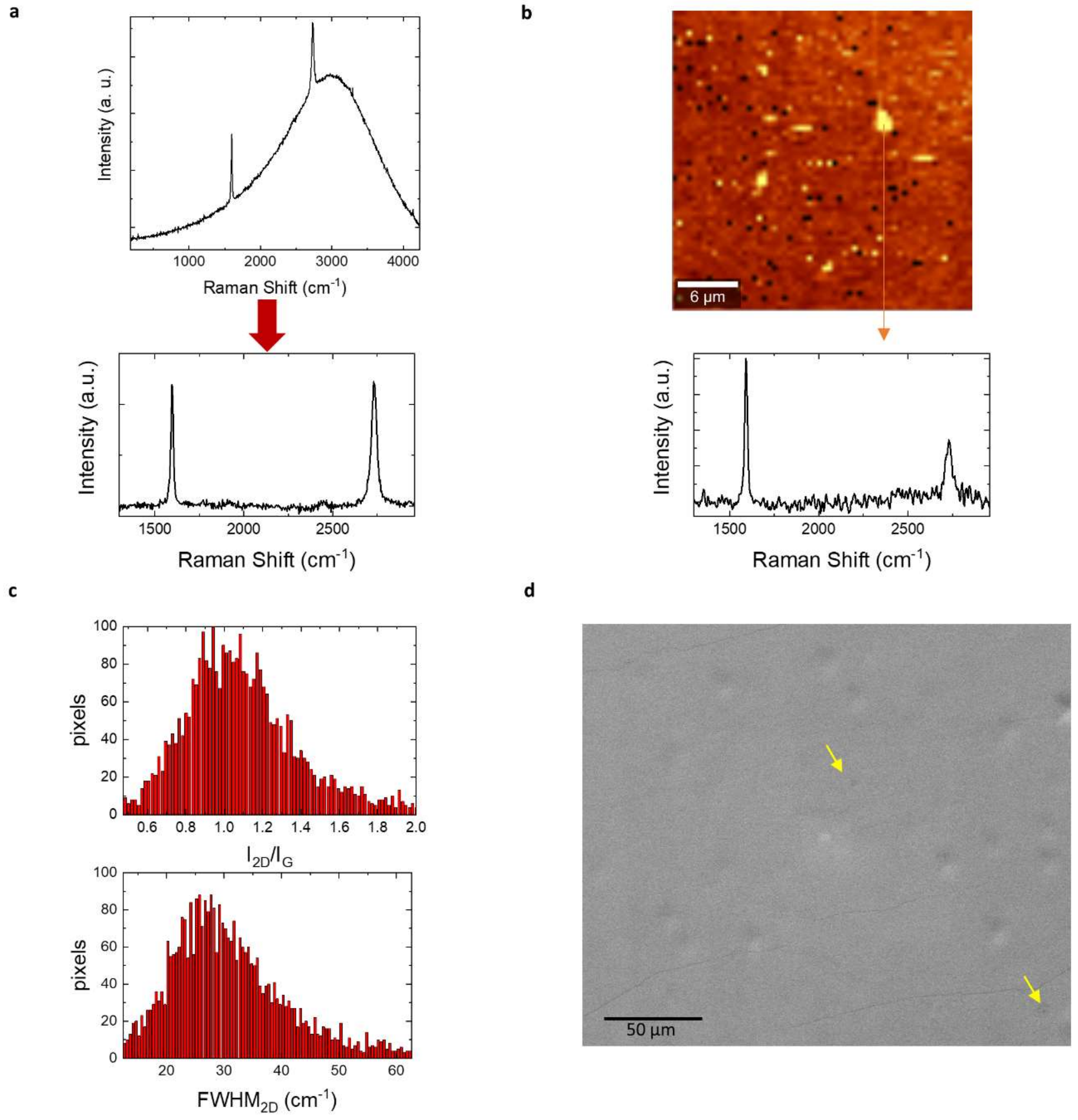


**Fig. S1 Characterization of chemical vapour deposition (CVD) grown graphene**. **a**, Raman spectrum of as grown graphene on copper. The absence of the *D* Raman peak, activated with disorder, indicates the good quality of the grown samples. Below, a Raman spectrum after copper background subtraction is shown; the two main Raman contributions are observed: *G* (1596 $cm^{-1}$) and *2D* (2730 $cm^{-1}$). **b**, Raman map of the *G* peak intensity where regions of higher intensity (yellow) correspond to second nucleation centers with characteristic Raman spectrum like the one shown below. **c**, Histograms of the *2D*/*G* intensity ratio and *2D* band width. 92% of the scan area (30x30$\mu m^2$) presents an intensity ratio of 2.4 characteristic of monolayer graphene. The average FWHM of the monolayer region is about 28 $cm^{-1}$. Regions of less than 4 $\mu m^2$ that show larger 2D FWHM correspond to the thicker (multilayer) nucleation spots. **d**, Scanning electron microscopy image of as-grown CVD graphene on copper foil; two second nucleation spots can be distinguished.

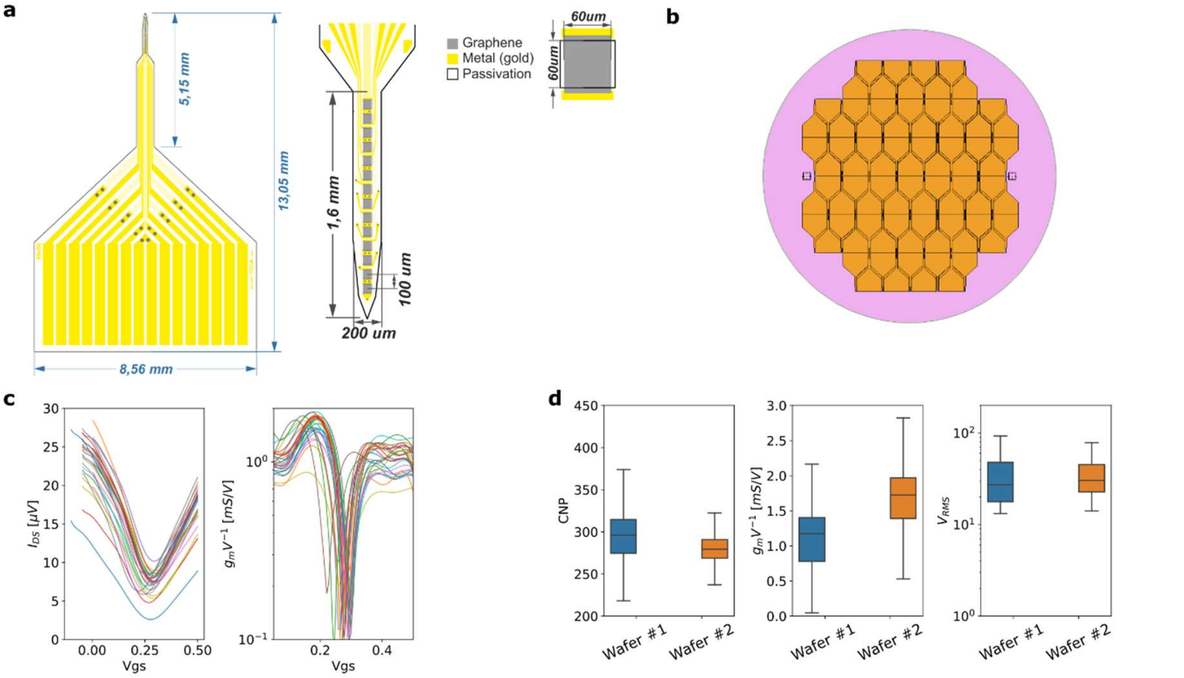


**Fig. S2 Design and electrical characterisations of the gDNPs. a**, Layout specification of the gDNP. **b**, The flexible polyimide gDNPs are fabricated with standard clean-room facilities on a 4-inch $SiO_2$ sacrificial wafers, enabling large production of neural probes for each batch. **c**, Transfer curves of all characterized gDNPs on one wafer with the corresponding normalized transconductance ($g_m$/V). Each curve is the average of 14 transistors on a gDNP. **d**, Distribution of charge neutrality point values (CNP), $g_m$/V and $V_{RMS}$ (integration range 1-2000 Hz) of all fabricated and characterised gDNPs on two different wafers, showing homogeneity, high yield and reproducibility of the technology.

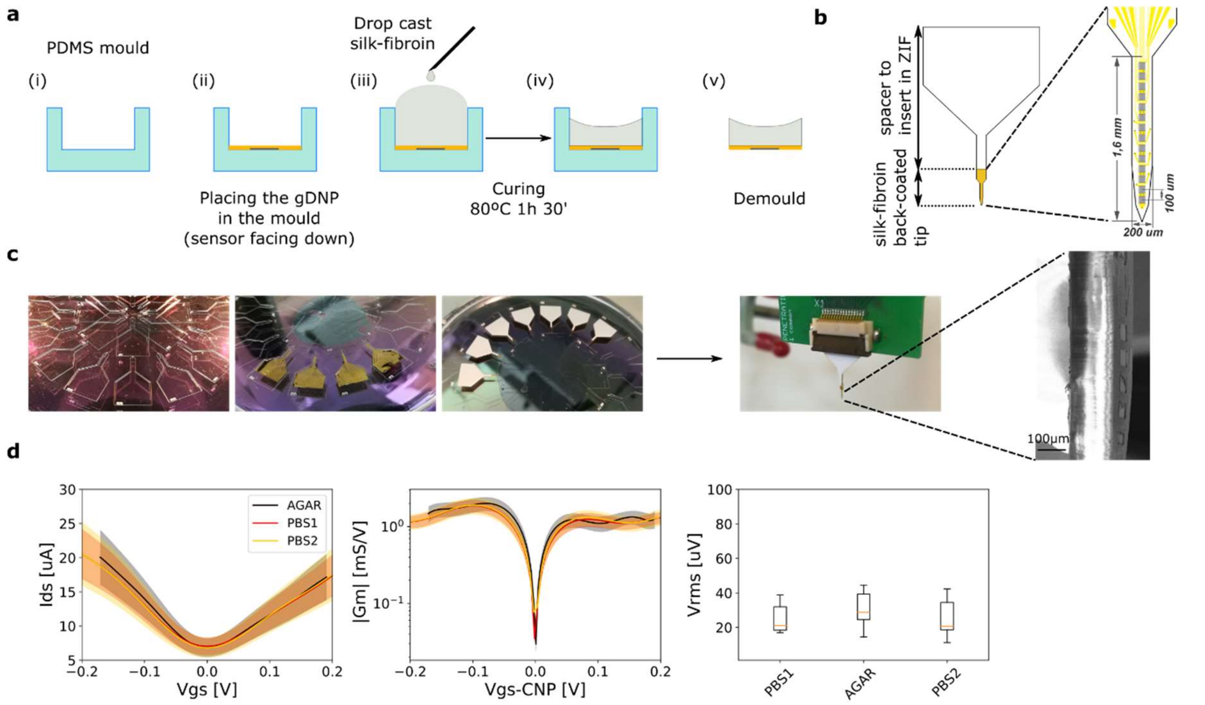


**Fig. S3 Back-coating of the gDNP with SF – functionality assessment**. **a**, Schematics of the back-coating stiffening technique: (i) cross-section of the PDMS mould; (ii) placement of the gDNP with sensors facing down-wards; (iii) drop-casting of the SF by filling the mould's trench. Double coat after 20min, then place on a hotplate and increase slowly the temperature to 80C; (iv) curing step of the SF at 80C for 1h and 30min; (v) remove from hotplate and gently demould the gDNP. **b**, gDNP design: a spacer (1mm thick) is attached to the back part of the gDNP in order to interface the ZIF and electronics. The tip of the gDNP is back-coated with SF. The very tip of the gDNP is the shank (width=200µm, length 1.6mm) with the 14 graphene transistors (100µm pitch). **c**, Pictures of the PDMS mould following the steps described in (**b**). After demoulding the back-coated gDNP, the probe is placed in the ZIF of the custom PCB to interface the electronics. Blow up of the tip: SEM image of the 150um SF-coated gDNP. **d**, Functionality assessment: Averaged transfer curve (Ids vs Vgs) and transconductance (gm vs Vgs) of a gDNP characterized before back-coating in 10mM Phosphate-buffered saline (PBS1); then after coating with SF and inserted in an agarose gel brain model (AGAR); After removal from agar, in 10mM PBS (PBS2). The stiffening protocol does not affect the performance of the gDNP.

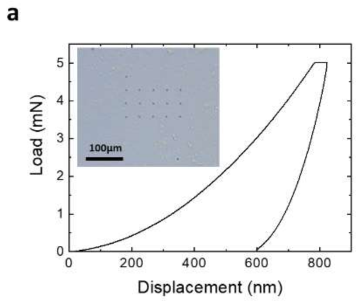


| SF cured@80ºC | | |
|---|---|---|
| Sample | H(MPa) | Er(GPa) |
| SF 1 | 369±7 | 10.7±0.4 |
| SF 2 | 365±2 | 10.1±0.1 |

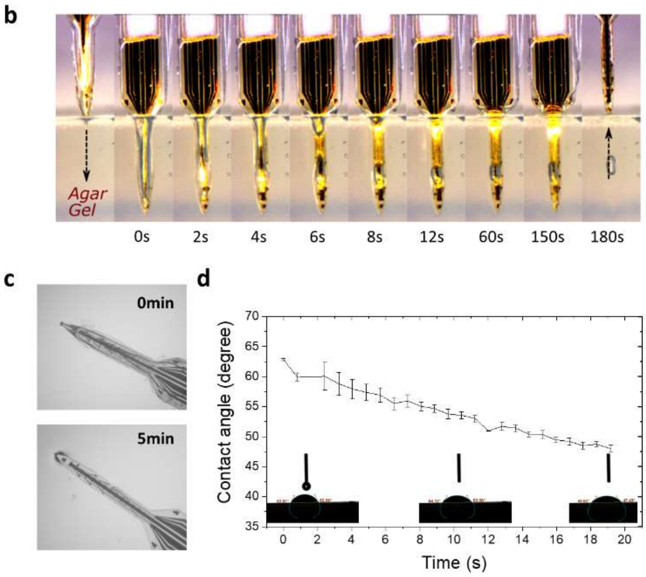


**Fig. S4 a** Representative indentation curve from the series of measurements performed to extract the hardness (H) and reduced Young's modulus ($E_r$) of silk-fibroin cured at 80ºC for 1h 30min on a $SiO_2$ chip (2cm x 2cm). Inset: microscope image of the indentation sites on the SF sample. The table shows the extracted values of **H** and **Er** as an average value of twenty indentations, performed on top of each sample. **b**, Image sequence of a SF coated gDNP at different time points during degradation in 0.6% agarose gel brain model. **c**, images of a SF-coated gDNP during degradation in agar just after insertion and after 5min. Most of the SF coatings are degraded within five minutes and delaminated from the gDNP surface. However, as silk fibroin is not water soluble, the degraded products often stays in the solution as small residue parts for a longer period of time. **d**, The hydrophilic nature of the SF was evaluated using contact angle measurements. As shown, the contact angle value is less than 90°, which implies that water can sufficiently wet the SF surface. Moreover, the contact angle value varied from 63° to 48° within 20s time, which shows the increasing absorption of water on the silk fibroin surface with time. SF coated probes showed good mechanical stability for their insertion for the first time. However, the coating material absorbs water from the surround quickly and loses its mechanical stiffness. Therefore, silk fibroin coated gDNPs are only suitable for single time insertion.

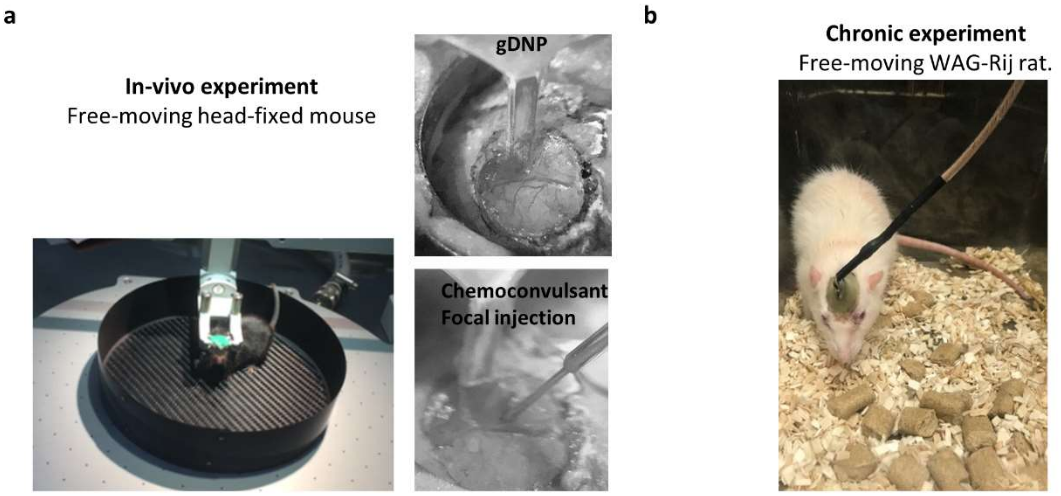


**Fig. S5 Pictures of in-vivo acute and chronic experiments. a,** Photographs of the free-moving head-fixed mouse undergoing habituation training, as well as craniotomy, gDNP and drug-injection positioning. **b,** Photographs of the WAG-Rij rat and the chronic implanted gDNP connected via a rotor to the electronics, enabling movement during the recordings.

| Experiment # | Neural probes | Animal model | *in-vivo* experiment | Chemoconvulsant | Sampling rate |
|---|---|---|---|---|---|
| 1 | gDNP (S35) | wild-type C57BL | Acute<br>Awake head-fixed ♦ | PTX-10 mM -250nL @ 50nL/s | 9.6 kHz/s |
| 2 | gDNP (S28) | wild-type C57BL | Acute<br>Awake head-fixed ♦ | 4-AP-50mM 200nL @ 20nL/s | 9.6 kHz/s |
| 3 | gDNP (S29) | wild-type C57BL | Acute<br>Awake head-fixed ♦ | 4-AP-50mM 200nL @ 20nL/s | 19 kHz/s |
| 4 | gDNP (S39) | wild-type C57BL | Acute<br>Awake head-fixed ♦ | 4-AP-50mM 200nL @ 20nL/s | 19 kHz/s |
| 5 | gDNP (S11) | wild-type C57BL | Acute<br>Awake head-fixed ♦ | 4-AP -50 mM -350nL @ 35nL/s | 9.6 kHz/s |
| 6 | gDNP (S15) | wild-type C57BL | Acute<br>Awake head-fixed ♦ | 4-AP-50mM 200nL @ 20nL/s | 9.6 kHz/s |
| 7 | gDNP (S26) | WAG Rij –absence epilepsy | Chronic<br>Freely moving ◊ | - | 9.6 kHz/s |
| 8 | gDNP (S4) | WAG Rij –absence epilepsy | Chronic<br>Freely moving ◊ | - | 9.6 kHz/s |
| 9 | gDNP (S32) | WAG Rij –absence epilepsy | Chronic<br>Freely moving ◊ | - | 9.6 kHz/s |
| 10 | gDNP (S22) | WAG Rij –absence epilepsy | Chronic<br>Freely moving ◊ | - | 9.6 kHz/s |

♦ Awake head-fixed in Neurotar frame with air-pump supported carbon-fibre frame
◊ Animals were freely moving and tethered without the use of anaesthesia. Connction via commutator.

**Table 1:** Overview of *in-vivo* experiments performed and used in this study to show the gDNP full bandwidth electrophysiological capability. All flexible gDNP were successfully implanted with silk-fibroin back-coating strategy as described in **Methods**.

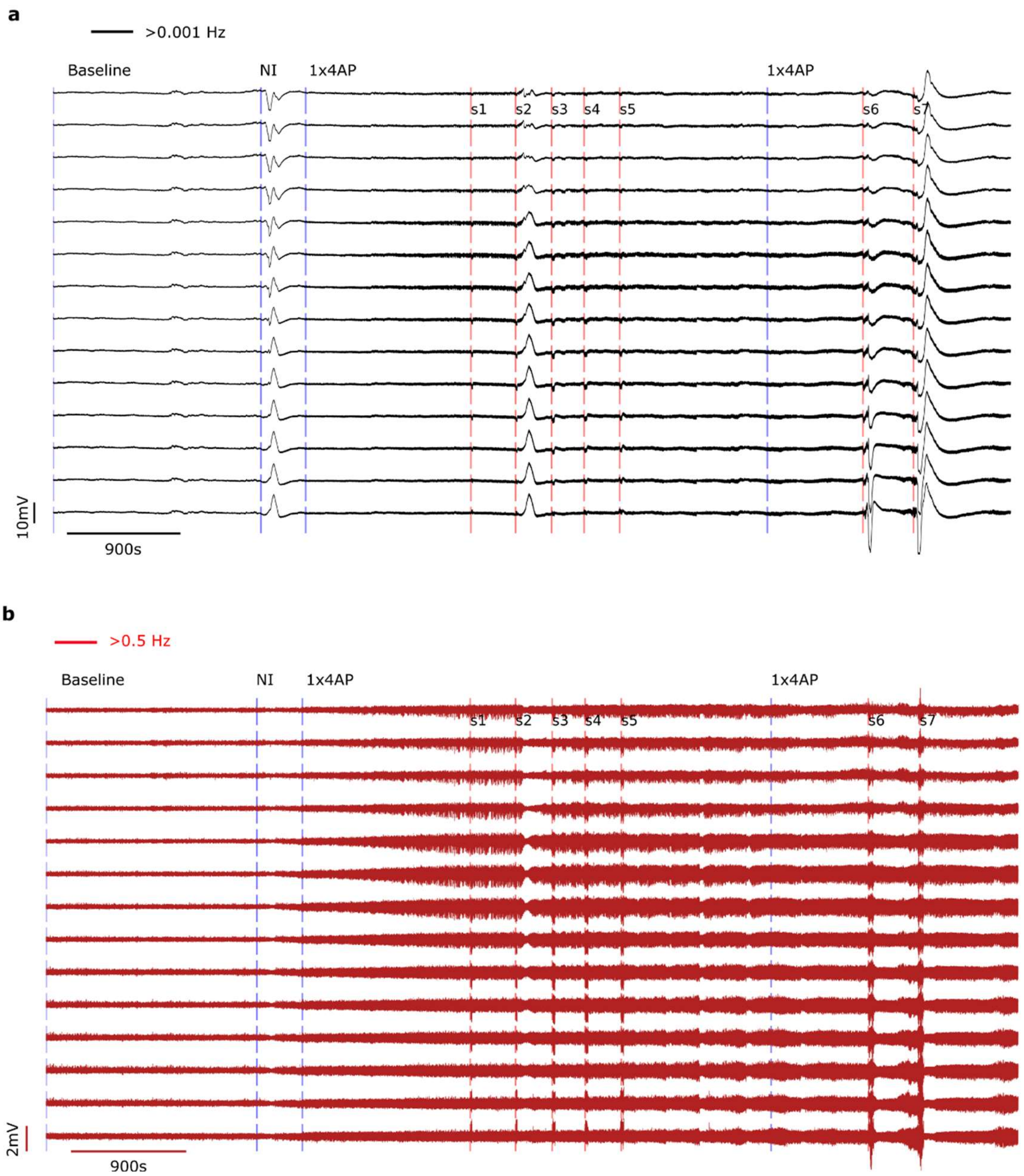


**Fig. S6 Long recording shown in Fig.2c for all channels on the gDNP**. **a**, Full-bandwidth recording filtered (>0.001 Hz). **b**, Same recording filtered above 0.5 Hz, showing the loss of low frequency information for conventional electrodes. Red vertical lines correspond to the seizure events (s1-s7) blue line describe the action during the in-vivo experiment, such as needle insertion (NI) and the injection of 4AP to induce epileptiform activity.

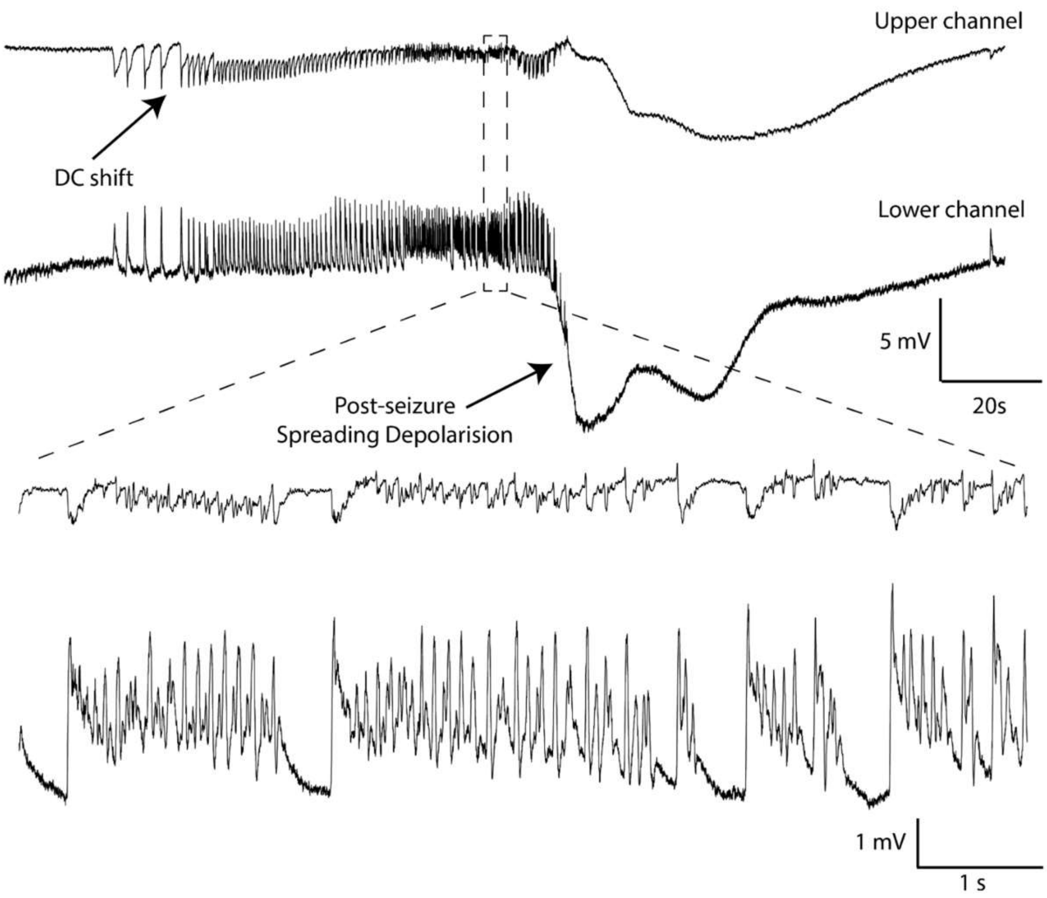


**Fig. S7 Example of seizure recorded with the gDNP in full bandwidth mode**. Upper and lower channels recordings, showing pre-ictal DC-shift, seizure and post-seizure spreading depolarisation. Blow up during the seizure illustrates the inversion in polarity as well as the amplitude differences between upper and lower channels (corresponding to neurons in the V1 cortex and CA1 region in the hippocampus, respectively).

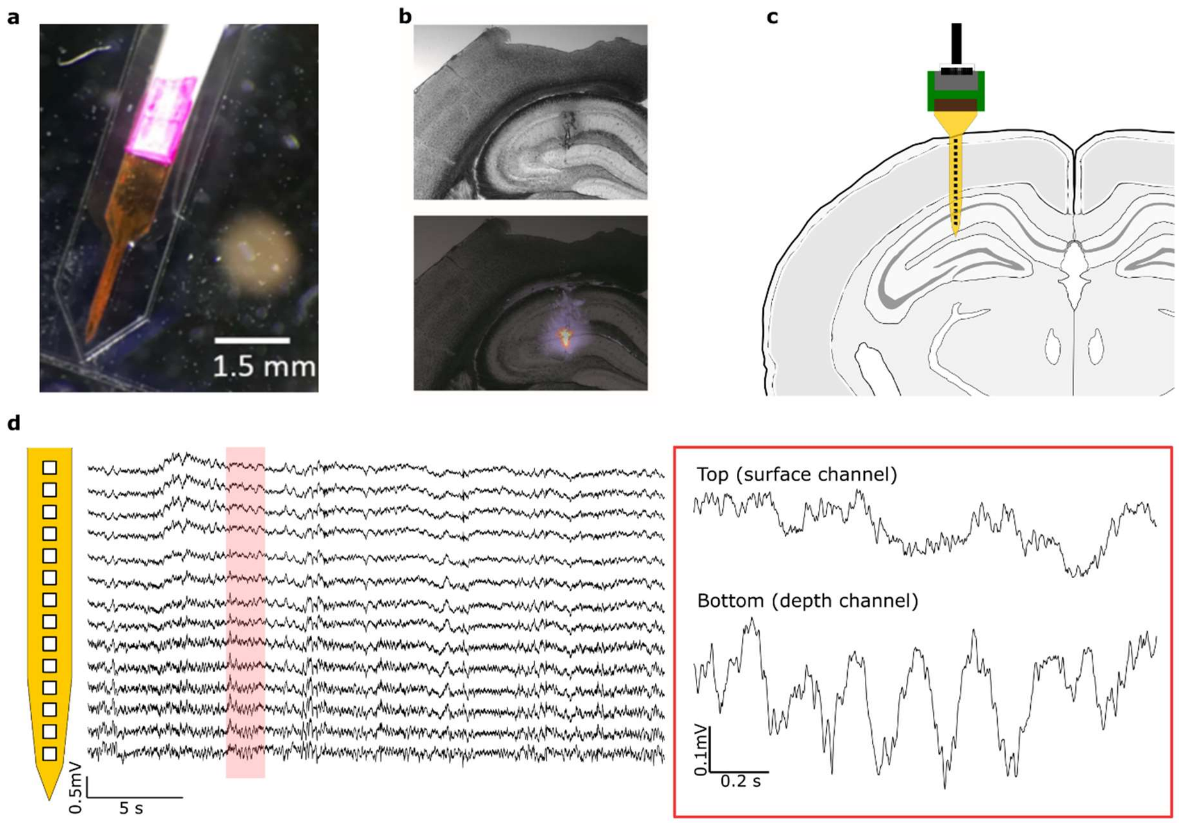


**Fig. S8 position of the gDNP in the mouse brain**. **a**, Picture of a gDNP in the PDMS mould before back coating with SF. The tip is coated with DiI stain before the SF is applied. **b**, Histology image of a coronal cross section of the mouse brain after the recording. DiI stain marks the position of the tip of the probe in the hippocampus. **c**, schematic of the position of the gDNP corresponding to the histology picture. **d**, Probe shape with profile recording during baseline activity (prior to drug injection) showing theta activity when the animal was moving in the lower channels of the shank, confirming that the lowest channels reached the hippocampus.

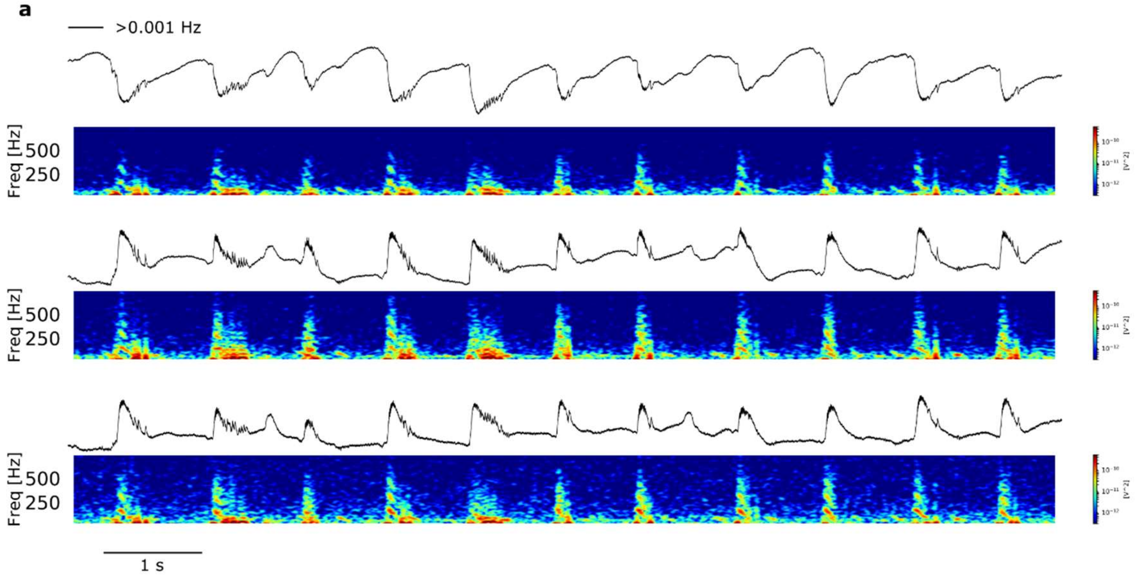


**Fig. S9 Interictal activity and HFOs.** Spectrograms (range 10-750 Hz) of the interictal activity with associated HFOs and sharp ripple waves in three different depths (shown in Fig3a). The spectrogram revels layer-specific high frequency tones of the HFOs. Top channel: superficial cortex; middle channel: lower layer of neocortex; bottom channel: CA1 region in hippocampus.

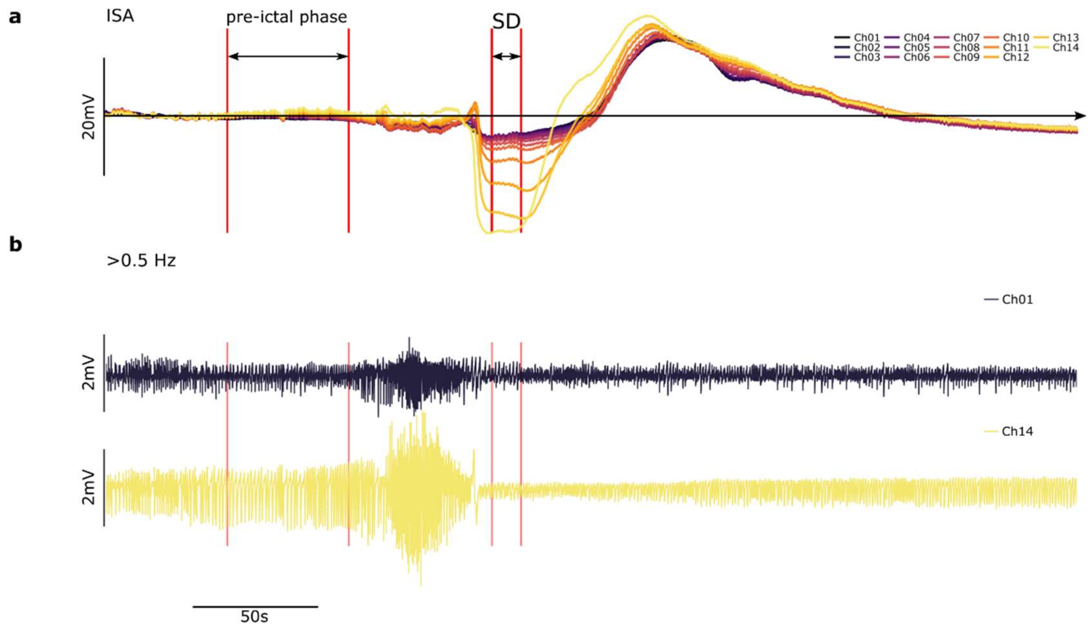


**Fig. S10 Description of the neural activity variation [%] and SD amplitude evaluation shown in Fig. 3d**. **a**, shows the LP filter (<0.5 Hz) of the recording shown in Fig. 3d. All the channels are overlapped with colour gradient corresponding to their depth. **b**, HP filtered (>0.5 Hz) recording for Ch01 and Ch14. The neural activity variation in Fig. 3d displays the neural silencing scenario that is happening after the hippocampal SD. The neural activity variation is evaluated in terms of ratio of the mean values of the RMS (sliding window of 1s) of the HP filtered (>0.5 Hz) signal during the SD and during the pre-ictal phase respectively. For the calculation of the amplitude of the SD and hyperpolarisation the minimum and maximum values shown in panel (**a**) are averaged in a short time window (2s). The amplitudes are calculated in reference to the zero set by averaging the first 50s of the pre-ictal phase.

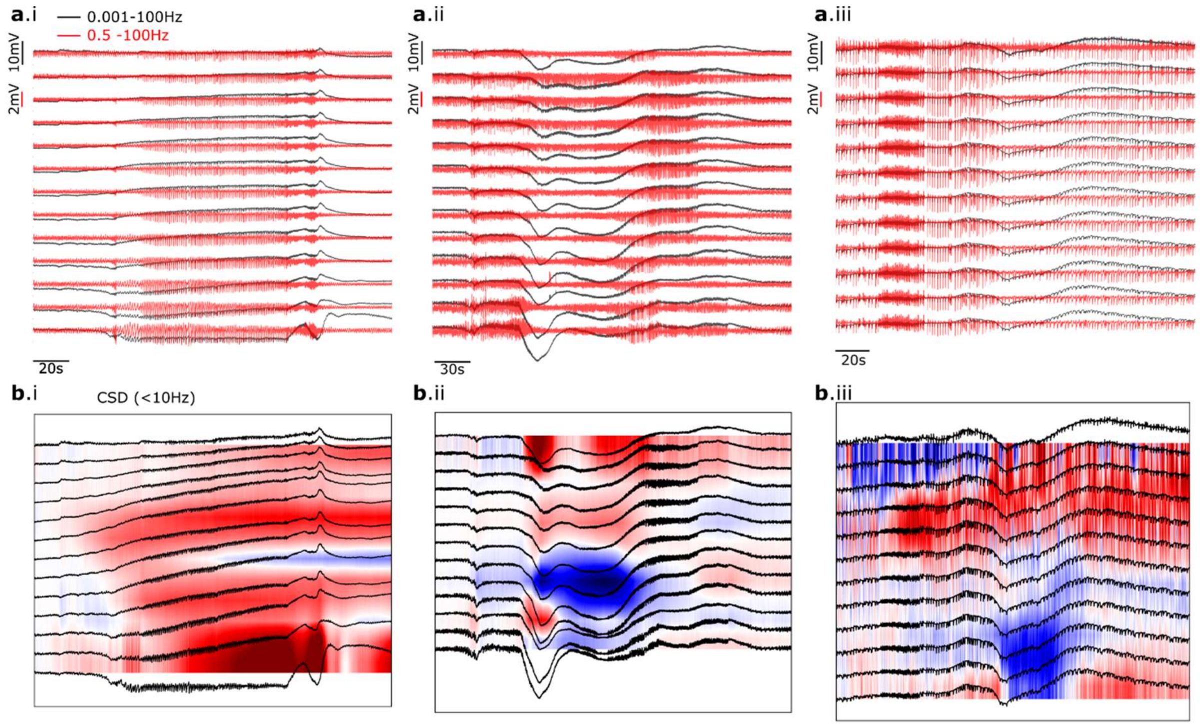


**Fig. S11 Current source density analysis of low frequency**. **a.**i-iii, profile recording during seizure events in three different mice showing seizure and spreading depolarisation. In black the signal is HP >0.01 Hz allowing detection of the ISA oscillations otherwise hidden to conventional electrodes due to HP filtering (red >0. 5Hz). **a**.i-ii, 4-AP was injected to elicit epileptiform activity; **a**.iii PTX was used as chemoconvulsant to induce epileptiform activity. **b**.i-iii, Current source density analysis (CSD) of the low-passed filtered full bandwidth recording (<10Hz) shown in (**a**) depicting different ionic dynamism during seizure in the three different *in-vivo* experiments. Usually the CSD analysis is applied to low-frequency part of the potential. The places where net current is entering or leaving the cell are called source (i.e. cations flowing from the intracellular space to the extracellular space) and sinks (i.e. cations flowing into the cell)[1,2].

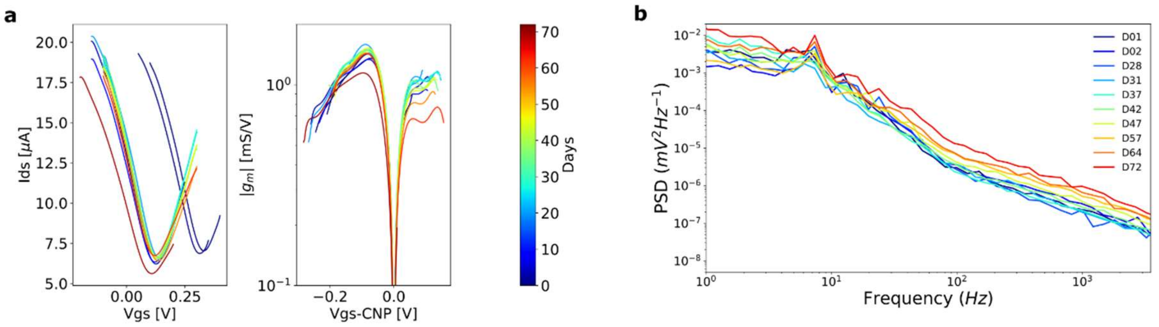


**Fig. S12 Electronic characterisation of the chronic-implanted gDNP in a WAG-Rij rat for over 10 weeks**. **a**, Averaged transfer curves and normalized transconductance ($g_m/V_{DS}$) performed in the rat brain of all gSGFET on the gDNP, showing stability of the device over 10 weeks. **b**, Averaged power spectral density (PSD) of all working channels on the gDNP over implantation time. The averaged PSD values are calculated over 500s of neural recordings. The peak around 8 Hz represents the frequency of the spontaneous SWDs.

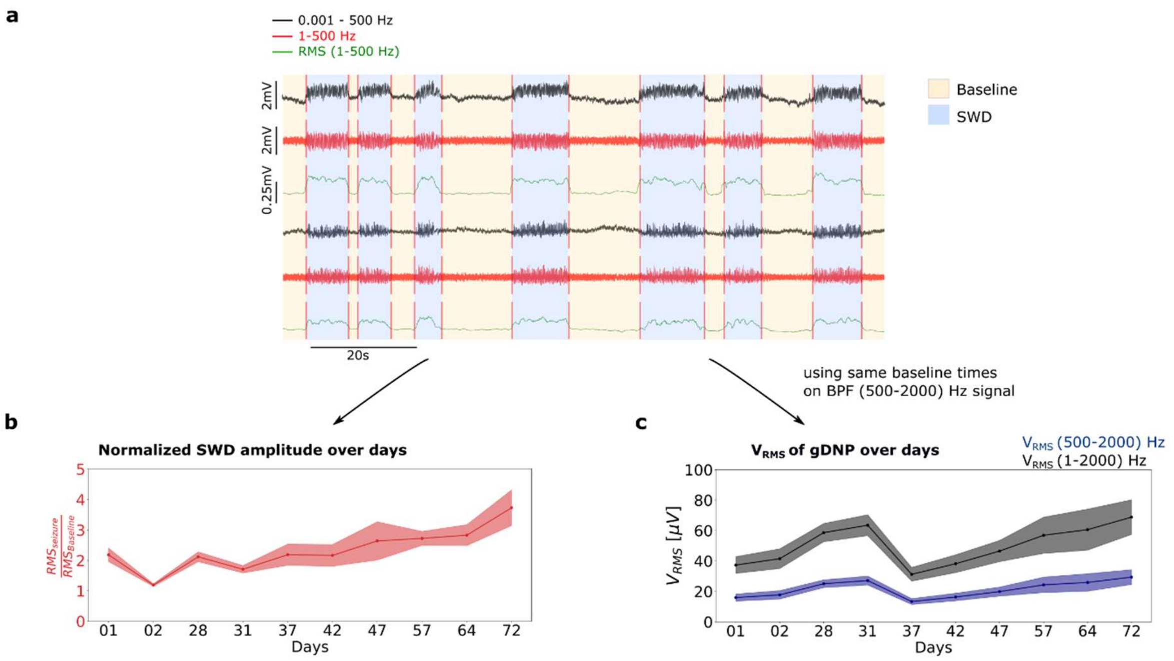


**Fig. S13 Description of the noise evaluation for the chronic experiment** (**Fig.4d**). **a**, The automatic detection of the SWDs is performed calculating the RMS (green) of the HP filtered signal 1-500 Hz (red) (sliding window 0.01 s). The mean value of the RMS along the whole recording is used as threshold level: everything above is considered SWD, everything below is baseline. **b**, normalised SWD amplitude over the implanted time. The ratio of the averaged RMS value during the SWDs and RMS value in the baseline is averaged for all channel on the gDNP. **c**, For the $V_{RMS}$ evaluation of the gDNP, we use the same detection strategy to filter out the SWDs event and average the RMS only in the baseline for the HP 500-2000 Hz. Then using the 1/f type of noise of graphene transistors[3] (see equation (1)), we extrapolate the $V_{RMS}$ of the gDNP in the range (1-2000) Hz using equation (2).

$$V_{RMS}^2 = \int_{f_1}^{f_2} S_V \, df = \int_{f_1}^{f_2} \frac{1}{f} \, df = ln\left|\frac{f_2}{f_1}\right| \tag{1}$$

$$V_{RMS,BW2}^2 = V_{RMS,BW1}^2 \frac{\ln(\frac{f_{f,BW2}}{f_{i,BW2}})}{\ln(\frac{f_{f,BW1}}{f_{i,BW1}})} \tag{2}$$

Where Sv is the noise power (1/f type of noise), $f_{f,BW2}$ , $f_{i,BW2}$ are the final and initial frequencies of the bandwidth BW2 and $f_{f,BW1}$ , $f_{i,BW1}$ the initial and final frequencies of BW1. In our case BW2=[1-2000Hz] and BW1=[500-2000 Hz].

**a**

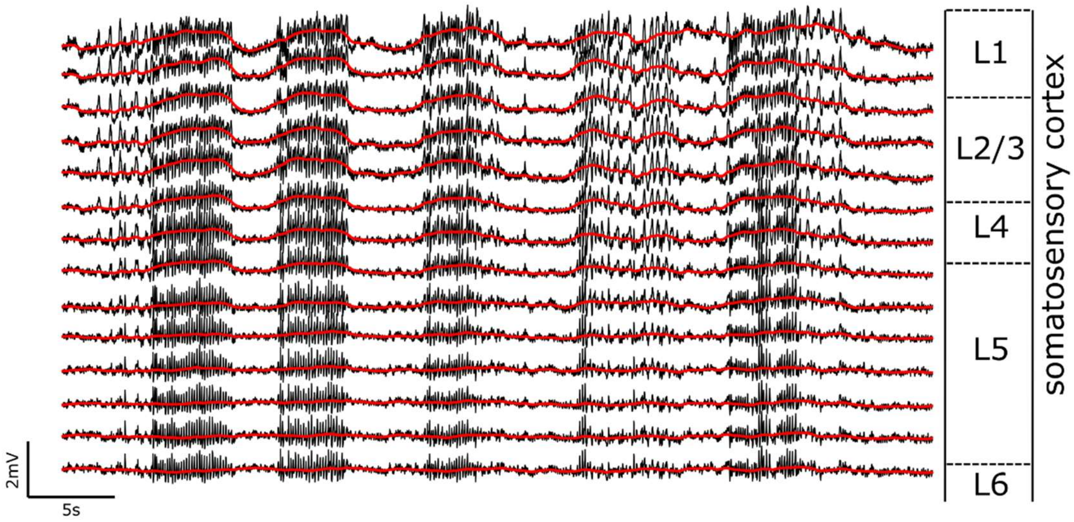


**Fig.S 14 DC-shifts during SWDs in WAG Rij**. **a,** Profile recording across the cortical laminae, illustrating SWDs through cortex layers. Overlapped low frequency component (< 0.5 Hz red) of the signal showing a gradient in amplitude along the rat's cortex. Largest DC shifts are observed in the more superficial layers.

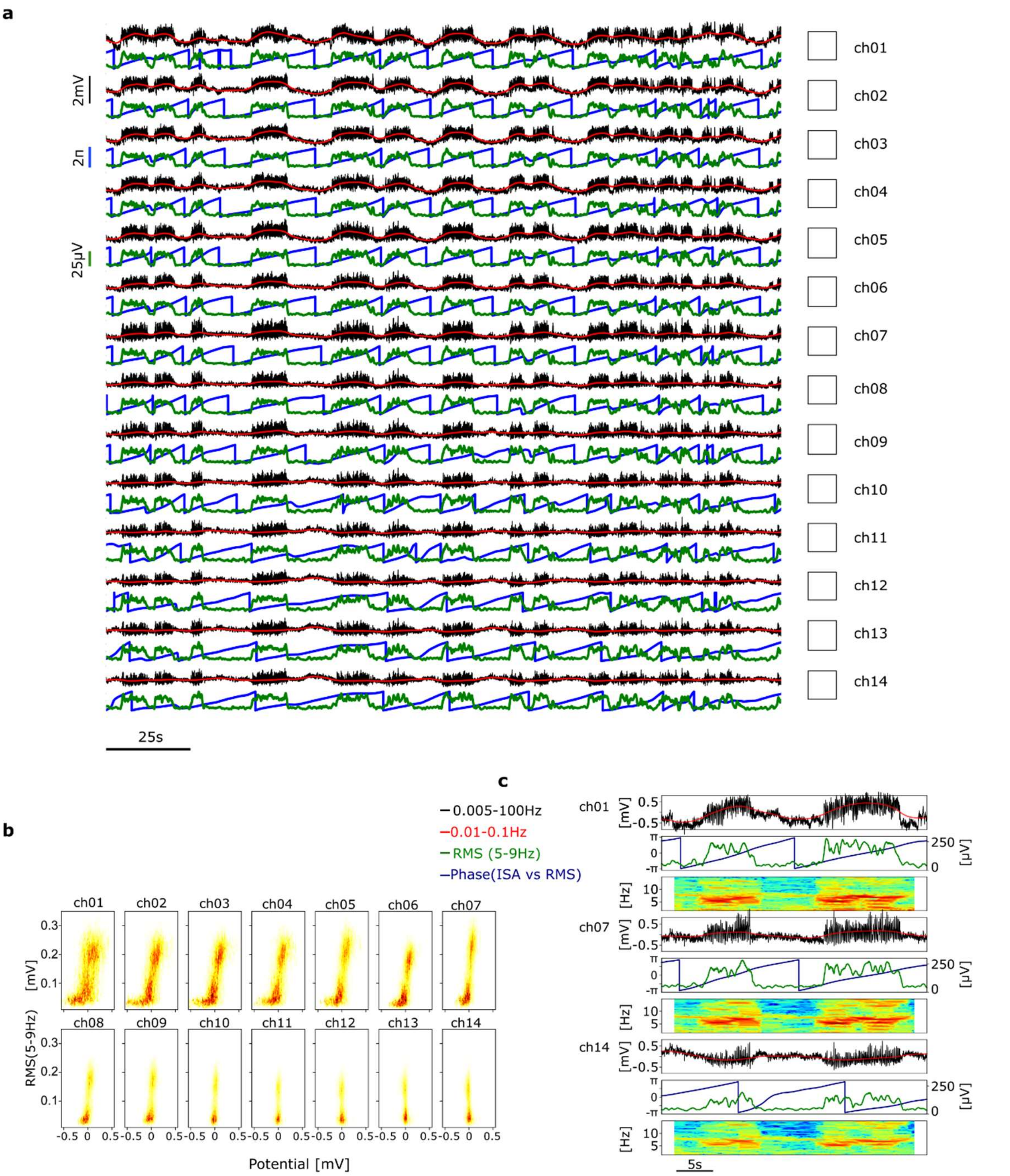


**Fig.S 15 a,** Vertical recording profile across the cortical laminae (somatosensory cortex) of the WAG-Rij rat illustrating the layer-specific DC-shift during the absence seizure (SWD). Root-mean square (RMS, green) calculated between 5-9 Hz and ISA phase (0.01-0.1 Hz blue). **b,** Histograms for all channels of the gDNP of the RMS (5-9 Hz) with ISA amplitude (BPF: 0.01-0.1 Hz) deflection calculated over 1600s of recording. **c**, Blow-up of three channels along the gDNP (Ch01, Ch07, Ch14) showing how the phase dependence of the ISA with the SWD changes with depth in the rat cortex.

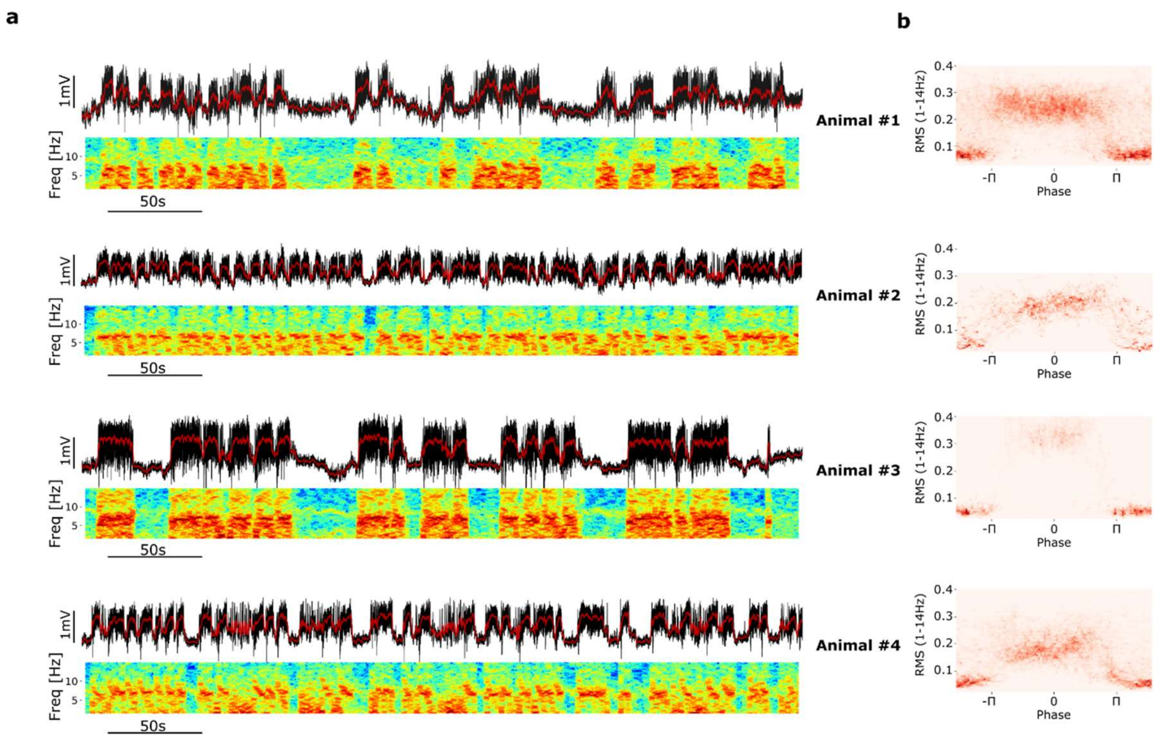


**Fig. S16 a,** Long recording (380s) showing the relationship between ISA and SWDs observed in all four implanted WAG-Rij rats. The full bandwidth (black) and HP <0.5 Hz (red) are overlapped. Spectrograms of the activity (1-14 Hz) to visualize the recruitment of SWDs. **b**, Density distribution for each WAG-Rij rat evaluated over a long recording (1600s). y-axes correspond to the RMS (1-14 Hz) associated with SWD while in the x-axis the phase of the ISA (0.005-0.05 Hz) computed by Hilbert transformation (red represents higher density).

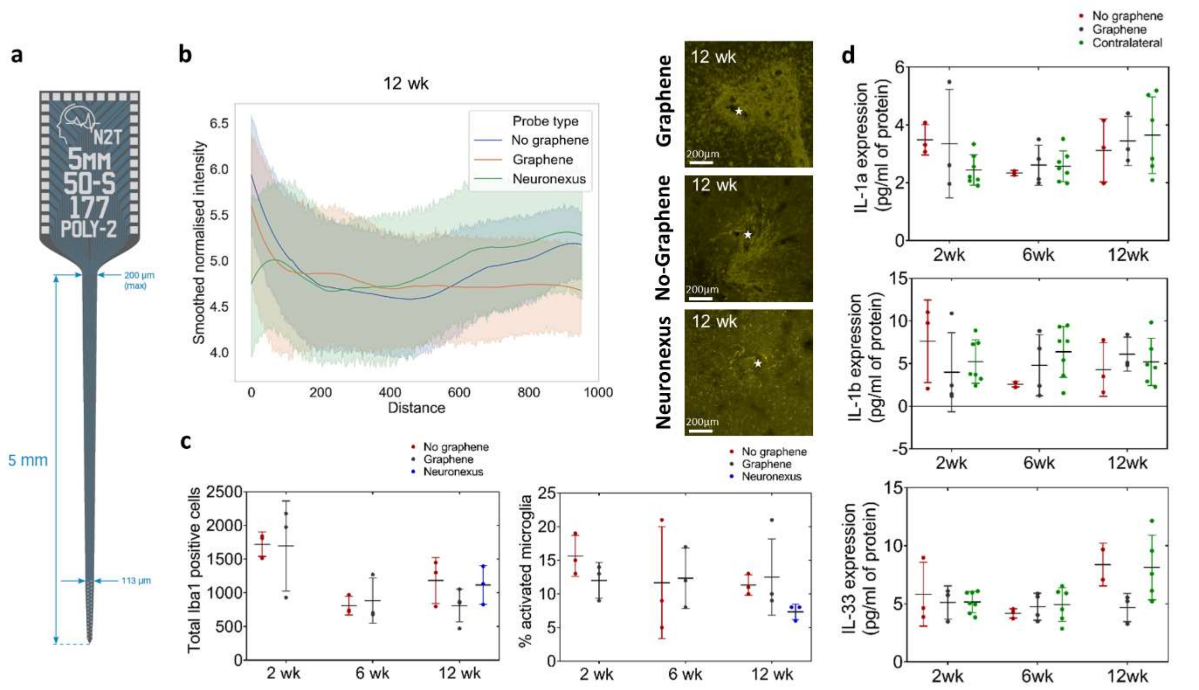


**Fig. S17 Immunohistochemical analysis a**, Neuronexus probe design (Source: Neuronexus product description). **b**, 12 week GFAP fluorescence intensity in the area surrounding probe implantation site for "no-graphene" gDNPs, "graphene" gDNPs and rigid Neuronexus probes. **c**, Microglial cell counts showed that there were a greater number of microglial cells present in the area surrounding device implantation at 2 weeks, but that this number subsided by 6 weeks. At 12 weeks, there was a similar number of microglial cells in the area for all devices implanted, however there was a slightly lower percentage of activated microglial cells in animals implanted with Neuronexus devices. **d**, ELISA data for pro-inflammatory markers showed no evidence of increased neuro-inflammation for either gDNP with and without graphene at any timepoint, even when compared to the contralateral hemisphere, where there was no device implanted.

To put this histological analysis in context, we also include to the analyses of Fig. 4 i-j the results of the cohort of animals implanted with rigid depth neural probes (Fig. S17c**,** NeuroNexus). Considering the narrower tip of the rigid probe compared to the gDNP design, no significant histological differences at 12-weeks post-implantation were observed.

The immunohistochemical analysis does not show any significant accumulation of pro-inflammatory cytokines; the upper graph in Fig. S17d, corresponding to the concentration of the cytokine IL-17a, suggests that there is no sign of an inflammatory response over the 12-week period for either device used, when comparing to the analysis of the contralateral hemisphere (where no probe was implanted).

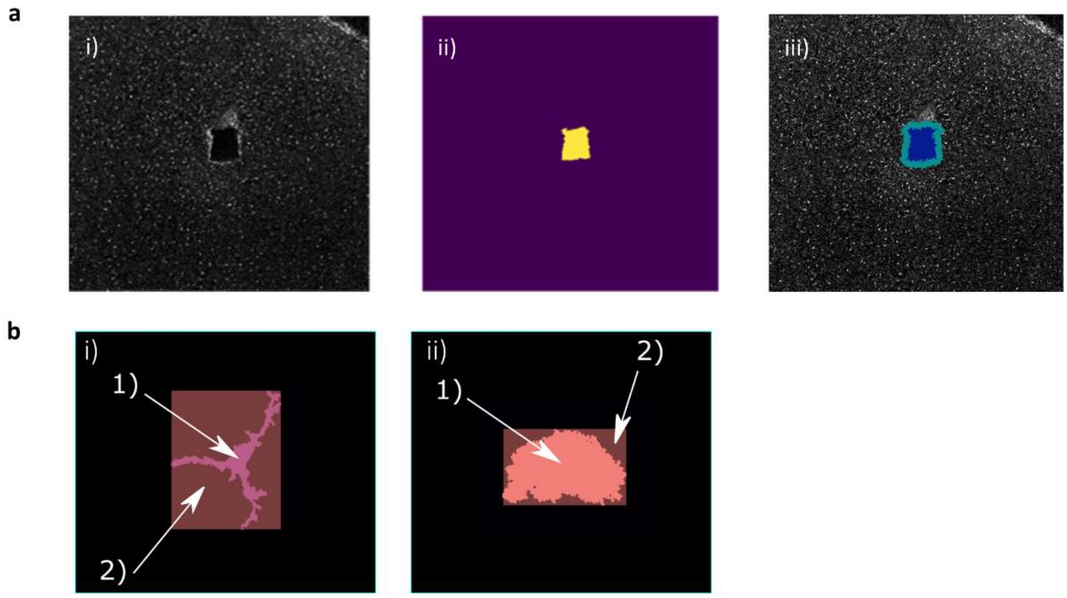


**Fig. S18 a**, Schematic diagram indicating strategy used for quantitative analysis of histology images. General strategy used for analysis of each type of fluorescence image: i) Original image; ii) Probe site located; iii) Having located the probe site, a binary dilation operation is applied to increase the size of the mask induced by the probe site by one pixel in every direction. The area and intensity in the outer band are calculated, then the process is repeated. This figure is for explanatory purposes only, and is not to scale. **b**, Quantifying microglia activation based on ratio of area of bounding box (2) to area of cell (1): i) Unactivated cells are small with long dendrites, so have a large bounding box. ii) Activated cells are rounder with no dendrites, so the cell body occupies most of the bounding box.
